# Recurrent evolution of cryptic triploids in cultivated enset increases yield

**DOI:** 10.1101/2025.02.14.638218

**Authors:** Yann Dussert, James S. Borrell, Jonathan Stocks, Harriet V. Hunt, Oliver W. White, Paul Wilkin, Richard Buggs, Lucie Büchi, Sebsebe Demissew, Feleke Woldeyes, Ilia J. Leitch, Wendawek M. Abebe, Richard A. Nichols

## Abstract

The high incidence of polyploidy in crops could be explained if domestication involved the selection of polyploids (cytotypes with more than two sets of chromosomes). Ancestral horticulture could have targeted desirable traits including yield, robustness to stress and to disease, rather than requiring knowledge of polyploidy itself. We find evidence for this process underway in enset (*Ensete ventricosum*, Musaceae) – a highly resilient crop constituting the main staple for over 20 million people in Ethiopia, which is clonally propagated and cultivated for its starch-rich corm and pseudostem. Prior to this study, enset was thought to be exclusively diploid (2n = 2x = 18). Using a newly-assembled chromosome-scale reference genome, and sequence data from 723 wild and domesticated enset individuals from southwestern Ethiopia, we demonstrate that around 20% of cultivated enset clones are triploid. We show that triploidy has arisen multiple times independently, that the triploid lineages have been given distinct landrace names, and planted disproportionately frequently. As well as providing evidence for enduring selection on triploids, our results reveal valuable genetic diversity captured by these triploid lines. They represent key resources for scientifically directed breeding of this major crop in the Ethiopian agrosystem, and could improve food security in sub-saharan Africa.

**Author Summary:** Duplication of a species entire genome (polyploidization) plays an important part in evolution. Domestication may have favoured polyploid crops including wheat, coffee, potato, cotton, strawberry, and banana. It has been proposed that this trend is explained by the selection of advantageous traits exhibited by polyploids compared to their relatives with unduplicated genomes. These desirable traits include bigger fruits, seeds or other edible organs, and better tolerance to stress and disease. We unexpectedly find that a large proportion of cultivated enset lines have three copies of their genome (triploidy) instead of two (diploid). Enset is a highly resilient crop in the banana family; its stem and leaves are processed to provide one of the main food sources for over 20 million people in Ethiopia. Prior to our study, enset was thought to be exclusively diploid. Farmers use landrace names that distinguish diploids from triploids without having this new genetic information, showing the differences between them are recognised and given importance. Triploids were planted disproportionately and appear to grow faster, suggesting they are being selected for their higher productivity. These triploid ensets are an important genetic resource for scientifically directed breeding of new varieties, which could improve food security in sub-saharan Africa.

## Introduction

Polyploidy, where an organism possesses more than two copies of its genome, has an important and pervasive role in plant evolution and speciation [1–3]. Polyploidy is a common feature of plant domestication, as crops have on average more often experienced polyploidization than their wild relatives, notably in monocots [4], sometimes after, but more commonly before domestication [4,5]. Polyploid crops often have higher productivity, often attributed to the larger size of cells and organs, or fixed heterozygosity due to the combination of divergent parental genomes, or masking of deleterious mutations [6]. Polyploids can display a higher tolerance to biotic and abiotic stress, such as increased resistance to disease or improved ecological tolerance [7,8], which might favor adaptation to the disturbed habitats that characterize cultivated landscapes. Finally, the increased sterility of crosses with diploids produces instantaneous reproductive isolation, facilitating fixation of domestication traits valued by the farmers for practical or even aesthetic reasons [9]. In general, polyploids with an even number of genome copies tend to be fertile, but polyploids with an odd number of genome copies tend to be infertile, as chromosome pairing fails at meiosis. Most polyploidy crops have an even number of genome copies, such as hexaploid bread wheat and octoploid strawberries. A well-known exception to this is the major commercial cultivar of banana (*Musa acuminata*), Cavendish, which is triploid and clonally propagated.

A closely related genus to bananas and plantains (*Musa*), *Ensete* (Musaceae) contains seven species of giant perennial herbs variously distributed across Africa and Southeast Asia [10]. Taxonomically split from *Musa* by their different chromosome base number (x=9 in *Ensete*, versus x=10 or x=11 in *Musa* [11]*), the seven published chromosome counts in Ensete* have indicated that the genus is wholly diploid. One species of *Ensete, E. ventricosum* (hereafter also referred to by its common name, enset), is a regionally significant domesticated crop in the Ethiopian Highlands, where it constitutes the main staple for over 20 million people, occurring ubiquitously across the south-western part of the country as the predominant starch crop [12]. Wild *E. ventricosum* occurs from South Africa to the Horn of Africa, but despite this broad distribution, has only ever been domesticated in Ethiopia, where circumstantial evidence suggests it has likely been cultivated for centuries or millennia [13,14]. Unlike bananas, domesticated enset is not cultivated for its fruit, but for its starchy pseudostem (trunk-like part of the plant) and underground corm, which are processed into a range of foods. Farmers clonally propagate preferred lines by removal of the pseudostem, a process which stimulates the spontaneous development of multiple shoots which are subsequently severed from the parent corm and transplanted [12]. Cultivated enset varieties are genetically differentiated from wild *E. ventricosum*, which co-occurs in the Ethiopian Highlands but reproduces sexually [15], typically by outcrossing. As enset is monocarpic (i.e., dies after seed production) and fruiting uses up stored resources, farmers rarely allow cultivated lines to flower and set seeds. However, this can happen in abandoned fields or if there is an over-abundance of food supplies.

Enset is a high-yielding and highly ecologically resilient and flexible crop, whose role in food security in Ethiopia earned it the epithet “the tree against hunger” [13]. Cultivated enset is grown in a wide range of environments across southern Ethiopia, over an altitudinal range of 1,500 to 3,000 m above sea level. Despite all this, enset is still an under-utilized and under-researched orphan crop which has the ecological and genetic potential to enhance food security more widely in sub-Saharan Africa. Recent years have therefore seen a number of genetic and genomic studies on enset and its wild relatives, including draft genomes assembled from short reads [15–19]. These studies did not challenge the assumption that *E. ventricosum* is diploid.

In this study, we assembled a chromosome-scale genome for enset using long read sequencing and genome conformation mapping, greatly improving available genomic resources for this species. By mapping reduced-representation sequencing data from 723 wild and cultivated individuals to this genome, we demonstrated the unexpected contemporary presence of both diploid and triploid enset in cultivation. We proceeded to analyse the geographical distribution and evolutionary relationship of diploid and triploid clonal lineages. We tested whether enset triploid individuals offered yield advantages to farmers and hence interrogated the dynamics of triploid clone selection in this close relative of banana.

## Results

### High proportion of triploid individuals in cultivated enset

As published assemblies for *E. ventricosum* were highly fragmented, we produced a new chromosome-scale reference genome for the species. Using PacBio HiFi and Omni-C sequencing, we sequenced enset leaf tissue grown from a seed of the cultivated variety *Maze*. The resulting pseudo-haploid assembly (806 contigs, total size: 534 Mb) had nine large scaffolds, ranging from 35.9 Mb to 68.0 Mb in size and containing 92% of the assembly (S1 Fig, S1 Table). Gene prediction, informed by RNA-seq data from four different tissues, produced a total of 35,238 protein-coding gene models. The assembly and gene prediction included respectively 98.6% and 96.2% of conserved BUSCO orthologs. A k-mer analysis (S2 Fig) showed that the sequenced individual was diploid and had high genome-wide heterozygosity (1.10%).

To investigate the genetic diversity and population structure of enset, we conducted reduced-representation genome sequencing on 658 cultivated individuals collected in south-western Ethiopia at multiple farms along altitudinal transects (T1 to T10), and 65 wild individuals from locations across the partly sympatric wild range (S3 Fig), mapping read data to our genome assembly. We assessed ploidy level by examination of allelic balance (AB, i.e. the proportion of reads supporting each major allele for heterozygous sites) in each individual (Fig 1A). Distributions for diploid individuals are expected to have a modal value around 0.5, and triploids around 0.66. Thirty individuals (4%) had aberrant distributions, generally with higher modal values (peak around 0.9 appearing to be an artifact of the cut-off parameters used for allele calling). Whole genome resequencing of six individuals followed by k-mer analysis (Fig 1B, S4 Fig) confirmed the diploid and triploid assignments. Resequenced aberrant individuals were either diploid or triploid, indicating that their AB data were unreliable, so all of them were excluded from subsequent analyses. All wild individuals were diploid. Among cultivated individuals, 414 were diploid and 214 were triploid. Individual inbreeding coefficients were generally highest for diploid plants and lowest for triploids (Fig 1A), meaning that triploids generally displayed a higher proportion of heterozygous sites. This lower individual inbreeding coefficient values of the triploid individuals suggests that they arose via outcrossing, not selfing.

**Figure 1.**
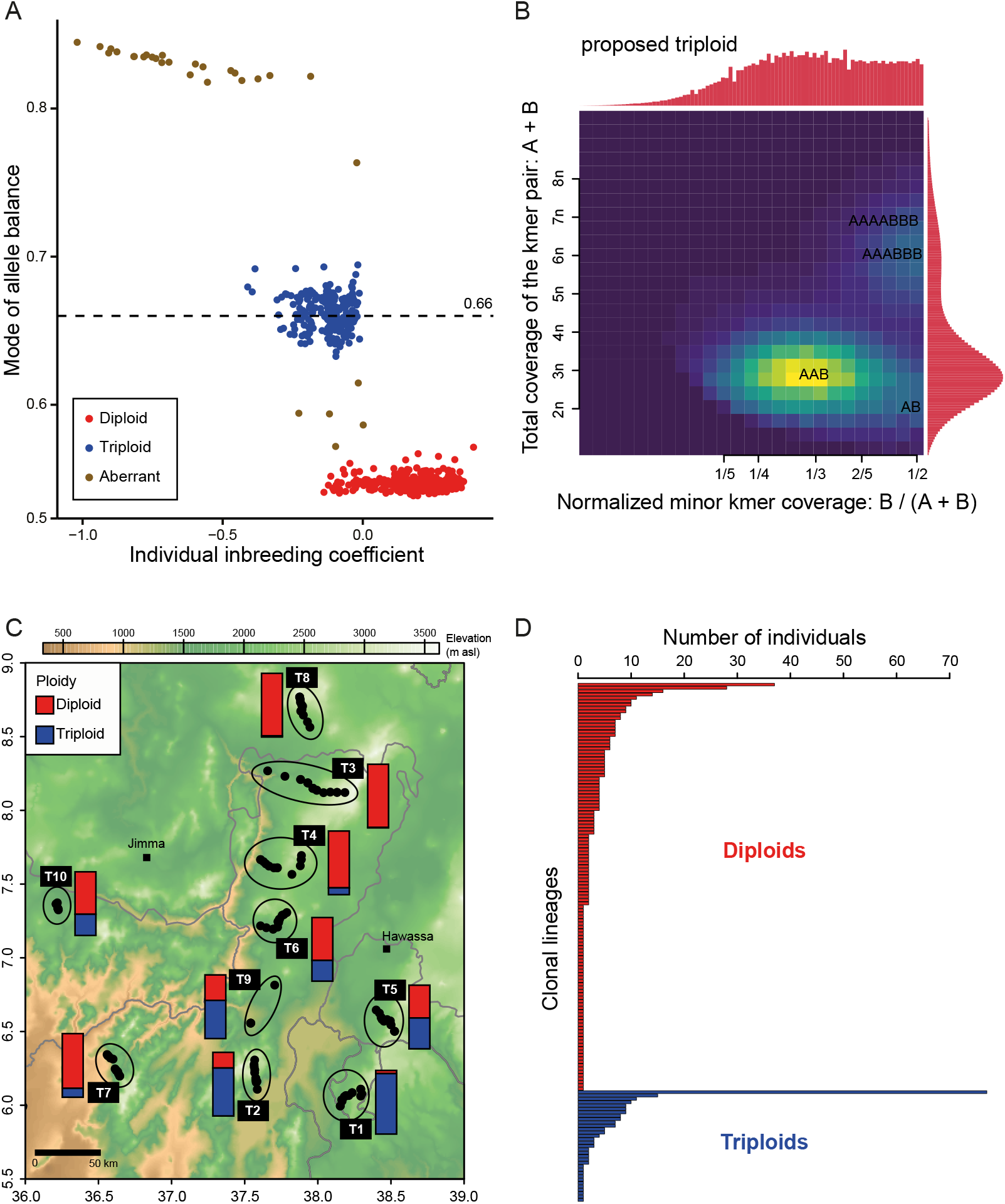
Ploidy variation in *Ensete ventricosum*. A: Individual inbreeding coefficient and mode of allele balance distributions for 658 cultivated and 65 wild individuals. Allele balance is defined as the number of reads supporting the most covered allele divided by the total number of reads for heterozygous genotypes. **B: K-mer profile of a triploid individual**. Heatmap colors indicate the number of heterozygous (i.e., differing by one nucleotide) 21-mer pairs in each bin, from 0 in dark purple to the maximum value in bright yellow. Histograms represent the total coverage of k-mer pairs for each axis. The brighter “smudge” indicates the ploidy of the sample, i.e. AAB for triploidy. **C: Geographic distribution of triploid individuals**. The proportion of cultivated individuals with a diploid or triploid cytotype is represented by a stacked barplot for each transect. Points on the map indicate sampling sites, and sites belonging to the same transect are encircled by an ellipse. Light gray lines delimit regional states. **D: Distribution of cultivated individuals in clonal lineages**. Clonal lineages, represented as bars, were determined on the basis of pairwise Euclidean genetic distances, separately for diploids (in red) and triploids (in blue).

Triploid cytotypes were more common in the southerly sites than the northerly ones (Fig 1C). A fitted multiple linear regression model (R^2^ = 0.56, F(3,79) = 35.39, *P*-value = 1e^-14^) revealed that the proportion of triploids per site, ranging from 95.1% for a site in T1 to 0.9% for a site in T3, was significantly correlated with latitude (*P*-value < 1e^-10^). There was no change in their proportion with altitude (S5 Fig, *P*-value = 0.31).

### Landrace names differentiate triploid from diploid enset

We identified clonal lineages among our samples using pairwise genetic distances (Fig 1D, S6 Fig). Within the 414 cultivated diploids there were 121 clonal lineages, of which 56 were represented by a single individual. Within the 214 triploids there were 33 clonal lineages, of which 11 were represented by a single individual. Seventy-seven (36%) of the triploids were from a single clone, suggesting preferential planting of this clone by farmers. By contrast, only 37 (9%) of diploids belonged to the most dominant diploid clone. After correcting the number of diploid clonal lineages by rarefaction (subsample size: 214), diploid ensets displayed a clonal diversity 2.6 times higher than the triploid group. We did not observe any clonal relationships among wild individuals, consistent with sexual reproduction in wild enset. Three cultivated diploid individuals (Ens_3135, Ens_3115 and Ens_3149) appeared to be genetically very close to wild individuals, suggesting occasional introductions of wild individuals into fields.

Farmers reported 177 vernacular landrace names for the sequenced cultivated plants, with 86 of them (around 49%) designating only one individual in our sample. To some extent the names identified specific clonal lineages (S7 Fig): 69.8% of pairwise comparisons of the same name were for the same clone. Farmers’ names consistently differentiated diploids from triploids: only 2.6% of pairwise comparisons of the same name were between individuals with different ploidy (compared to a random expectation of 14%, permutation test, P-value <10^-4^). Thus farmers unconsciously classify plants according to their ploidy based on phenotype alone. Farmers’ names differed with geography and ethnic group. For example, the most widespread triploid clone was commonly called *Ganticha* in T1 and T5 (48 samples), *Maze* in T2 (13 samples), *Wulanche* or *Wolanche* in T6 (3 samples), *Toracho* in T5 (3 samples), as well as ten other names applied to single individuals. This is unsurprising, given that at least 65 languages are spoken across the enset cultivating region.

### Multiple independent triploidization events

We found clear genetic differentiation between wild and domesticated enset, as described by previous studies [15,17,20,21]. This was seen in the principal component analysis (PCA, Fig 2A) and the clustering analysis (S8 Fig). The only exceptions were the three cultivated diploids that had already been found to be very closely related to wild samples based on genetic distances (see above), and two other cultivated diploid individuals, which were placed between domesticated and wild individuals in the PCA. These two individuals also showed a high proportion of membership of wild clusters in the clustering analysis (S8 Fig), indicating that they might have originated from recent sexual crosses between wild and domesticated individuals. These patterns suggest occasional introduction of wild individuals into fields by farmers, and rare gene flow occurring between the wild and domesticated groups. No evidence for such translocations or gene flow was found for triploids.

**Figure 2.**
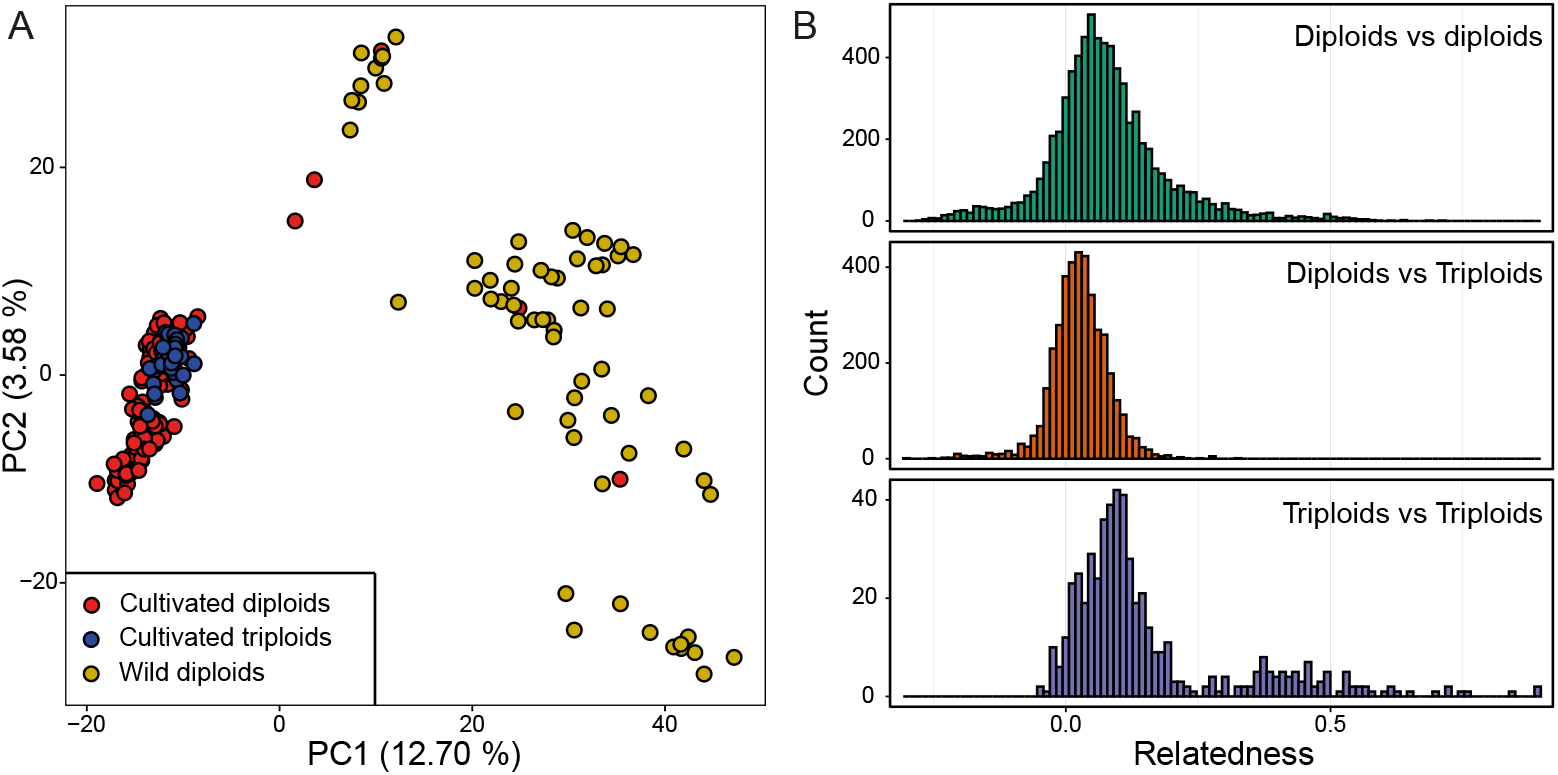
Population genetic structure and relatedness in *Ensete ventricosum*. **A: Principal component analysis on the clone-corrected dataset.** Diploid and triploid cultivated individuals are colored in red and blue respectively, and wild samples are in yellow. **B: Relatedness between pairs of cultivated individuals**, as measured by Huang method of moment estimator.

Both diploid and triploid cultivated enset were genetically differentiated from wild enset (diploids: F_ST_ = 0.198; 95% Confidence Interval: 0.190-0.208; triploids: F_ST_ = 0.209; 95% CI: 0.200-0.216). Differentiation was low between triploid and diploid cultivated clones (F_ST_ = 0.032, 95% CI: 0.030-0.034). In the PCA based on SNP data, triploids overlapped with a subset of the cultivated diploids (Fig 2A), and the clustering analysis did not group them separately (S8 Fig). This suggested that triploid samples originated from sexual crosses between diploid domesticated individuals, with no involvement of wild individuals. Analysis of kinship among samples showed that relatedness was low between most diploid-triploid pairs of lineages (Fig 2B, middle panel), suggesting that the direct diploid progenitors of extant triploid lineages are not found in our sample set. Triploidization may have occurred elsewhere, a substantial time ago or that the diploid progenitors are rare or extinct. Most of the triploid clonal lineages were not related to each other (Fig 2B, bottom panel), indicating that they probably had different diploid parents, though a few triploid lineages with high pairwise relatedness might have originated from one or two of the same parents. This suggests triploids have arisen from among a population of cultivated diploids through multiple independent events.

### Triploid ensets have higher pseudostem volumes than diploid ones

In addition to sequencing, we measured the pseudostem, one of the main plant parts harvested for food, of domesticated plants as a proxy for plant yield. Using a Bayesian linear mixed-effects model, we compared the pseudostem volume of diploids and triploids. We used ploidy as a fixed effect with age and bioclimatic variables (summarized by the first two principal components of a PCA) as covariates to the ploidy. Membership of clonal lineages and a genetic covariance matrix between lineages were applied as random effects (see Methods). The explanatory power of our model was high (conditional R^2^ = 0.64, 95% Credible Intervals: 0.60-0.68), and around 13% of the variation in pseudostem volume was explained by differences between clonal lineages (marginal R^2^ = 0.51, 95% CI: 0.39-0.60). We found no effect linked to environmental PC1, which was negatively correlated with altitude (Pearson’s *r* = -0.93). We found an interaction between age, ploidy and environmental PC2 (Fig 3, S2 Table), which was correlated with latitude (Pearson’s *r* = 0.76). While the pseudostem volume as a function of age varied for diploids along PC2, this was not the case for triploids. Overall, the estimated pseudostem volume of triploids was higher: for example, a triploid harvested at 6 years old would be 42 to 75% larger than a diploid, after controlling for other factors.

**Figure 3.**
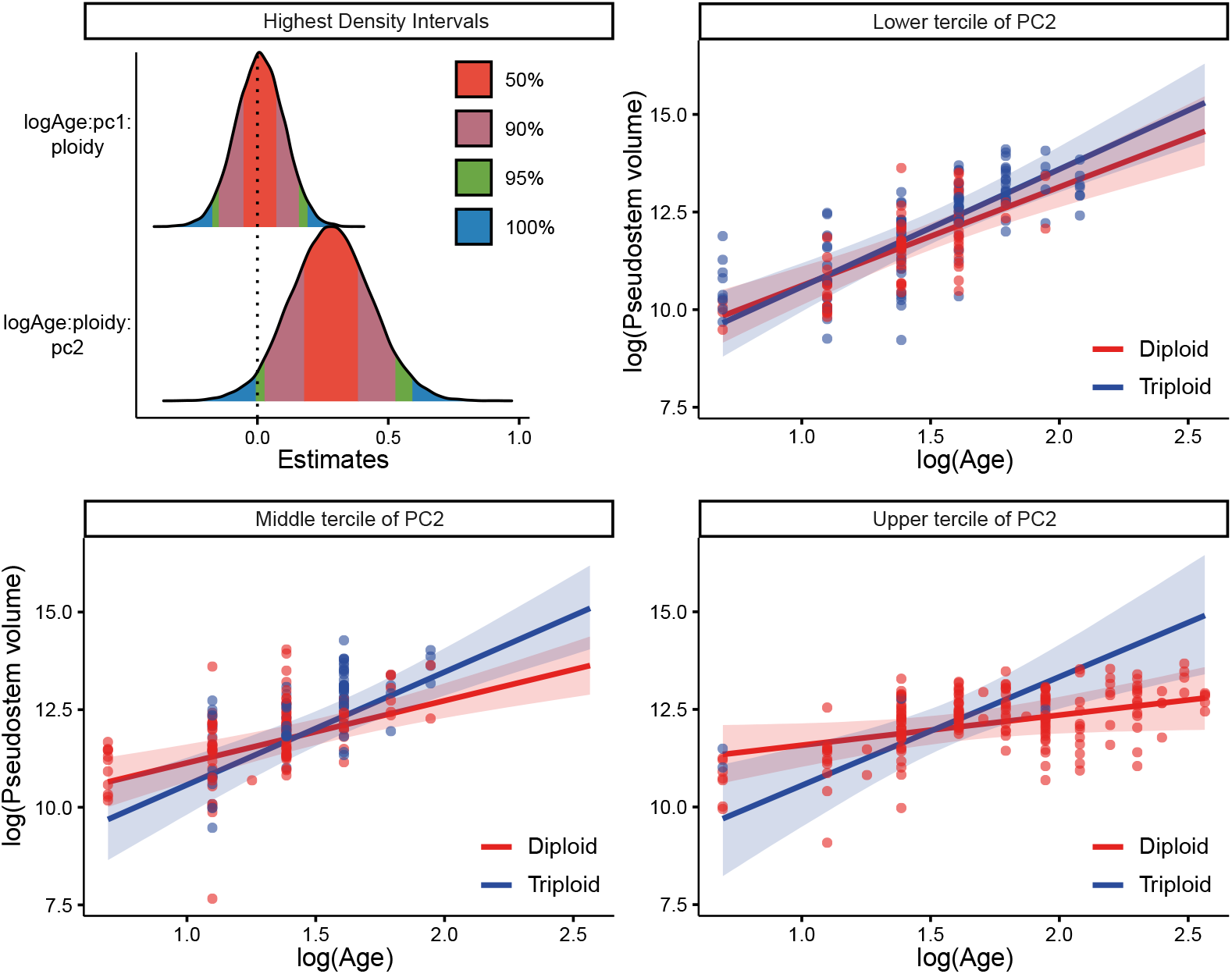
Bayesian linear mixed-effects analysis for pseudostem volume in cultivated diploid and triploid *Ensete ventricosum*. Left panel: estimation of fixed effects of the model. Effect size for ploidy is represented relative to the diploid category. Colored surfaces correspond to different values for the highest density interval of the full posterior distributions. Right panel: Interaction effect between plant age, ploidy, and PC2 (from a principal component analysis on environmental variables) on the plant pseudostem volume. Estimated linear relationships between the log-transformed age and log-transformed pseudostem volume are represented as lines, and their 95% credible intervals as colored areas, for diploid (in red) and triploid (in blue) individuals. Linear relationships for the three terciles of PC2 values are displayed.

As we sampled and measured plants *in situ*, there are possible confounding effects that we could not take into account, such as differences in agricultural practices between areas (e.g., harvesting of plants at different ages or stages of maturity, differences of propagation process [22] or manure application) or recent differences of weather between sites, which could have influenced plant growth. With our results showing the importance of triploid enset in southwestern Ethiopia, the phenotypic findings should be investigated in common garden experiments, to assess the reproducibility of the advantages of triploids that we detected across a range of environments.

## Discussion

For many polyploid crops, knowledge about the timing of polyploidization, and the number of parallel events involved, relies upon speculative reconstruction of events in the distant past. The discovery of mixed diploid and triploid lineages in enset, a regionally important staple, is an ideal opportunity to study the trajectory of polyploidization in relation to domestication in real time. Evidence of spontaneous and agronomically advantageous polyploidization in enset adds to a growing number of crops for which an increase of the number of chromosome sets has proved useful and been selected for by farmers [5]. Our results indicate that triploidization in enset was recurrent, and probably occurred after the main domestication period. Other examples of post-domestication autopolyploidization include yams [23,24], taro [25], or triploid apple cultivars [26]. Our results also present a correlation between clonality and polyploidy, as has been observed in other angiosperms [27]. In the case of the crops mentioned above and enset, their clonal agricultural propagation will have allowed them to avoid the costs of infertility resulting from triploidy.

As cell size is correlated with genome size, an increase in ploidy leads to bigger cells [28]. This, in turn, generally leads to an increase of the size of plant organs, as illustrated by a recent meta-analysis comparing autotetraploids and their diploid progenitors [29]. The pseudostem is one of the main organs harvested for food in enset, and its volume is correlated with the quantity of food products after processing [30]. Consequently, an increase in size would be an obvious advantage easily identified and selected for by farmers. Other important plant parts, such as the corm, which is also used for food, might also see a size increase, as its size is also correlated to the pseudostem diameter [31]. Besides an increase of plant organ sizes, changes in cell size also have important effects on plant physiology. In triploid citrus [32] or in tetraploid *A. thaliana* [33], *polyploidy has been associated with higher water stress tolerance for example. Increase of ploidy has also been linked with changes in cell wall composition in A. thaliana* [34], *which might be beneficial in enset for food processing. Genome size increase leads to higher costs in terms of nitrogen and phosphorus needs [35], but this might be alleviated by the fact that enset plots generally receive the largest quantity of inputs (i*.*e*., *manure) compared to other crops in Ethiopian farms. Apart from changes directly caused by cell size increases, a change of the number of chromosome copies also have important genetic effects. It can mask recessive deleterious mutations, but also beneficial ones, depending on their dominance level [6]. Phenotype changes may be mediated by dosage effects, e*.*g. if a triploid has two copies of an allele instead of one, as it has been shown in A. thaliana* and maize [36,37]. Future investigations of phenotypic and gene expression differences between triploid and diploid ensets will be necessary to understand the precise effect(s) and potential benefits of polyploidy in the species.

If triploid enset varieties display a higher yield and might also have other advantages compared to diploids, the existence of a latitudinal gradient for their presence is hard to explain. One possibility is that formation of triploid plants is more frequent in the south. Production of unreduced gametes has been linked to environmental conditions, notably high or low temperatures [38,39], and could also be associated with differences in farmer practices. Less intensive farming landscapes in the south [12] are likely to provide more opportunities for cultivated enset to flower, leading to subsequent outcrossing and hence an increased likelihood of polyploid formation. Triploids might also be better adapted to environmental conditions in the south. Finally, we might be observing an on-going process, where a few competitive triploid lineages, that originated in the south by chance, are selected by farmers and replacing diploid varieties and expanding their distribution to the northern part of Ethiopia. The fact that one particular lineage represents such a high proportion of triploid ensets across the study area would support this. This would also mean that advantages brought by triploidy in enset are likely to depend substantially on the alleles carried by the parents and their interactions.

While the possible replacement of diploid varieties by a lower number of higher-yielding triploid landraces might be advantageous for farmers in the short term, it might induce a decrease of genetic and variety diversity *in situ*, as only a small number of triploid lineages were dominant in the south. This could lead to the loss of potential adaptive or beneficial alleles that could be useful for farmers or in future breeding programs. It could also result in an increased vulnerability to diseases, as observed in bananas, whose cultivation relies mainly on a single dominant clone [40]. Clonal lineages of enset are also vulnerable to accumulation of deleterious somatic mutations [15]. As revealed in this study, a large number of cultivated enset individuals, including important varieties (e.g. *Ganticha, Ado, Maze, Midasho* or *Kitisho*), are triploid. In other species, triploids are generally less fertile. Due to irregularities during meiosis, triploid individuals often produce aneuploid gametes [41,42], which can lower pollen or embryo fertility, as in banana [43] or grapes [44]. As enset is propagated vegetatively by farmers, this should usually not be an issue. However, outcrossing will be likely needed to enable future adaptation to rapid environmental change [15]. Sexual reproductive capacity in enset has not been studied thoroughly, and until now the existence of triploid individuals was unknown. A recent study [45] reported high seed viability in a sample of domesticated ensets including named varieties found triploid in our study. However, their ploidy level had obviously not been determined. Consequently, the impact of polyploidy on enset fertility is very much in question, so it is essential to assess the ploidy level and the fertility of plants used in breeding programs. On the other hand, we suggest that farm productivity and hence food security could be enhanced by the increased use of triploid enset, with the potential of breeding novel triploids, and exploration of its adaptive range. However, care should be taken to preserve the genetic diversity of domesticated diploid populations as a safeguard for the future.

## Material and Methods

### Enset reference genome assembly and annotation

#### DNA & RNA extraction and library preparation

The individual used for the genome assembly was grown from a seed collected in Wolaita (6.9°N, 37.8°E, 2,120m a.s.l.). This seed originated from a sexual cross between a plant with the vernacular name *Maze* and an unknown parent. The latter was probably a cultivated individual, since, to our knowledge, wild plants do not occur in this region. DNA was extracted using a CTAB protocol, then quantified with a Qubit 2.0 Fluorometer (Life Technologies, Carlsbad, CA, USA). The PacBio SMRTbell library (∼20kb) for PacBio Sequel was constructed using SMRTbell Express Template Prep Kit 2.0 (PacBio, Menlo Park, CA, USA) using the manufacturer recommended protocol. The library was bound to polymerase using the Sequel II Binding Kit 2.0 (PacBio). Sequencing was performed on PacBio Sequel II 8M SMRT cells, generating 21.6 Gb of data. For the Dovetail Omni-C library, chromatin was fixed in place with formaldehyde in the nucleus and extracted. Fixed chromatin was digested with DNAse I, chromatin ends were repaired and ligated to a biotinylated bridge adapter followed by proximity ligation of adapter containing ends. After proximity ligation, crosslinks were reversed and the DNA purified. Purified DNA was treated to remove biotin that was not internal to ligated fragments. Sequencing libraries were generated using NEBNext Ultra enzymes and Illumina-compatible adapters. Biotin-containing fragments were isolated using streptavidin beads before PCR enrichment of each library. The library was sequenced on an Illumina HiSeqX platform to produce around 30x sequence coverage.

RNA was extracted from four different tissues: leaf, midrib, pseudostem and roots. Total RNA extraction was done using the QIAGEN RNeasy Plus Kit following manufacturer protocols. Total RNA was quantified using Qubit RNA Assay and TapeStation 4200. Prior to library preparation, DNase treatment followed by AMPure bead clean up and QIAGEN FastSelect HMR rRNA depletion were performed. Libraries were prepared with the NEBNext Ultra II RNA Library Prep Kit following manufacturer protocols, and libraries were run on the Illumina NovaSeq6000 platform in 2 × 150 bp configuration.

#### Genome assembly

PacBio CCS reads were assembled with Hifiasm 0.15.4-r347 [46] with default parameters. BLAST results of the Hifiasm assembly against the nt database were used as input for blobtools 1.1.1 [47] and scaffolds identified as possible contamination were removed from the assembly. Finally, purge_dups 1.2.5 [48] was used to remove haplotigs and contig overlaps. Scaffolding of the HiFi assembly was carried out with HiRise, a software pipeline designed specifically for using proximity ligation data to scaffold genome assemblies [49]. Omni-C reads were aligned to the draft input assembly using bwa [50]. The separations of Omni-C read pairs mapped within draft scaffolds (mapping quality > 50) were analyzed by HiRise to produce a likelihood model for genomic distance between read pairs, and the model was used to identify and break putative misjoins, score prospective joins, and make joins above a threshold.

A chloroplast sequence was assembled with NOVOPlasty 4.2 [51] using short reads from another sequenced landrace published previously (GenBank BioProject PRJNA252658) [19]. This sequence was used as a query to search for chloroplastic scaffolds in our assembly with BLAST 2.10.1+ [52], keeping only hits with a length > 500 bp. We then computed the proportion of each scaffold covered by chloroplastic sequences using the coverage command in BEDTools 2.29.2 [53]. Scaffolds with > 80% BLAST hit coverage were discarded from the assembly (500 scaffolds for a total of 19.6 Mb). We also tried to assemble a mitochondrial genome using the same method, but without success. Consequently, in order to detect and discard mitochondrial sequences, we used mitochondrial gene sequences from *E. glaucum* [54] *as queries for BLAST searches, keeping only genes with a minimum query coverage of 80%. Scaffolds with a valid hit (with the exception of the nine largest scaffolds) were considered as potentially mitochondrial and discarded (80 scaffolds for a total of 5*.*4 Mb)*.

#### Repeat and gene annotation

A *de novo* library of repeated sequences was generated using RepeatModeler 2.0.3 [55] with LTR structural analysis enabled. The genome sequence was then annotated and softmasked using this library with RepeatMasker 4.1.2 [56]. Both RepeatModeler and RepeatMasker were used through the Dfam TE Tools container v1.5 (https://github.com/Dfam-consortium/TETools). We searched for canonical telomeric repeats (CCCTAAA / TTTAGGG) in 1 Mb windows along the genome sequence using the nuc command in BEDTools. To detect potential centromeric regions, we looked for large tandem repeat arrays in the nine largest scaffolds using repaver (available at https://gitlab.com/gringer/bioinfscripts). Centromeres in *E. glaucum* contain large arrays of the Egcen repeat and are enriched for an interspersed nuclear element (LINE) called Nanica [57], discovered initially in the centromeres of *Musa acuminata* [58]. *To confirm our initial annotation of centromeres, the sequences of Egcen and Nanica were used as queries for a BLAST search in our assembly, and their number per 1 Mb computed with BEDTools coverage. The results of these analyses were plotted using RIdeogram [59] in R. RNA-seq reads were trimmed using Trimmomatic 0*.*36 [60] (parameters: ILLUMINACLIP:TruSeq3-PE*.*fa:2:30:10:5:True LEADING:3 TRAILING:3*

*SLIDINGWINDOW:4:15 MINLEN:36) then mapped onto the new reference sequence with STAR 2*.*7*.*9a [61], keeping only properly paired reads with SAMtools 1*.*9 [62]. For protein evidence, we downloaded all Embryophyte protein sequences from the OrthoDB v10 database [63], and protein sequences from M. balbisiana, M. itinerans, M. schizocarpa* and *E. glaucum* from the Banana Genome Hub website [64] (https://banana-genome-hub.southgreen.fr). RNA-seq and protein evidence were both used to annotate protein-coding genes with BRAKER 2.1.6 [65,66] on the softmasked genome sequence.

#### Quality control

Quality control was carried out through comparison of the k-mer content between the PacBio CCS reads and the final genome assembly, with KAT 2.4.2 [67] using a k-mer length of 21. The k-mer spectrum for the PacBio reads was also analyzed using GenomeScope 2.0 [68]. The level of completeness of the genome assembly and the gene annotation was assessed using BUSCO 5.3.0 [69] with the embryophyta_odb10 database.

### Sampling of enset in southwestern Ethiopia

Enset individuals were systematically sampled across eight transects (T1 to T8), 20-50 km long, across the enset growing region (Supplementary Fig. 3). These transects were oriented along environmental gradients, generally encompassing >1,000 m elevational range and therefore capturing a broad range of local climates including temperature and precipitation variation. This approach sought to capture a substantial portion of genotypic diversity and locally adapted landraces, while minimizing geographic distance within transects and associated isolation by distance effects. For each transect we sampled 10 locations, collecting tissue from all landraces present across three farms. We generally sampled a 10 × 10 cm piece of cigar leaf, and immediately dried it on silica gel. We interviewed farmers to obtain the variety name and the age of collected individuals. Additionally, for each plant we measured the pseudostem (height, basal and apical circumferences), one of the principal harvested tissues for consumption. We also used a few accessions from two other different areas (T9 and T10), collected opportunistically. Cultivated accessions were supplemented with wild individuals. These predominantly occur to the west and north of the main cultivated enset distribution with a patchy occurrence. We sought to sample as comprehensively as possible across this range. Environmental covariates for sampling locations were sourced at a 1-km resolution from Chelsa v2.1 [70,71] and sampled using the Raster package in R software.

### Reduced-representation sequencing for population genomics

After freezing leaf samples in liquid nitrogen, DNA was extracted using the DNeasy Plant Mini Kit or the DNeasy 96 Plant Kit (Qiagen) and eluted in 70 μl of AE buffer. Quantification of genomic DNA was performed using a Quantus fluorometer (Promega). Samples were sent to Eurofin Genomics, where they were sequenced with Genotyping by Random Amplicon Sequencing-Direct (GRAS-Di), a reduced representation sequencing method using random primers [72].

Adapters were trimmed from reads using Trimmomatic, and read pairs with at least one read < 36 bp were discarded. Reads were then mapped onto our new genome assembly with bwa mem 0.7.17, keeping only properly paired reads with SAMtools. Small variants were called using freebayes 1.3.1 [73] (parameters: –min-mapping-quality 10 –min-base-quality 15 –no-population-priors), keeping only biallelic SNPs with a quality score > 20, less than 25% missing data and a minor allele count of 3. Low quality samples with > 45% missing data were discarded. We then discarded sites with a mean read coverage > 500 and a F_IS_ value > 0.8. Finally, we filtered out spurious SNPs potentially caused by copy number variation by detecting sites with a median allele balance (number of reads supporting the alternate allele divided by the total number of reads, for heterozygous genotypes) < 0.35 or > 0.65 and with no homozygous genotypes and sites with a proportion of heterozygous genotypes > 60%. These SNPs and any site closer than 1,000 bp from them were discarded. The final dataset included 20,173 SNPs.

### Identification of triploids

Individual inbreeding coefficients were computed with VCFtools 0.1.17 [74]. The distribution of allele balance (defined here as the number of reads supporting the most covered allele divided by the total number of reads, considering only heterozygous genotypes) for each individual was determined with vcfR [75] for sites with a read coverage > 20, and the mode of these distributions was computed with modeest [76] in R. We expected diploid accessions to have a mode around 0.5, since heterozygous sites should show a 1:1 allele coverage ratio. Values of 0.66 (1:2 ratio) would be associated with triploidy, and so on. Plotting of individual inbreeding coefficients against the allele balance modes for all samples revealed the existence of three groups (see Fig. 1A in the Results section). After visual inspection of the raw allele balance distributions, we classified individuals as diploid, triploid or aberrant.

To confirm the ploidy results found with the GRAS-Di data, we randomly chose one diploid, three triploid and two aberrant samples and resequenced their whole genome on an Illumina NovaSeq sequencer (2 × 150 bp paired-end reads). After adapter trimming with Trimmomatic, the k-mer spectrum for each individual was determined with Jellyfish 2.2.6 [77], and their ploidy was further explored with Smudgeplot 0.2.5 [68], using a k-mer size of 21. Additionally, reads were mapped onto the reference sequence as described above, and SNPs discovered with the GRAS-di data were genotyped again with freebayes to check the allele balance distributions for the resequencing data. Due to the lack of availability of a flow cytometer in Ethiopia, and restrictions on export of living material, we were unable to estimate ploidy level from genome size measurements by flow cytometry using fresh plant material. While we tested this approach by rehydrating silica dried samples, as used successfully for some plant materials [78], we were unsuccessful, most likely because samples were too old (2-3 years).

To ascertain whether ploidy level was correlated with vernacular landrace names, we calculated the proportion of pairs of individuals sharing the same name that also shared the sample ploidy: *P*_*p*_ .The observed value of *P*_*p*_ was compared with the distribution generated by randomly relocating the landrace names to individuals (10,000 iterations).

### Population genetic analysis of mixed ploidy dataset

For analyses of datasets with mixed ploidy, we followed previous recommendations [79]. First, SNP calling was carried out again for triploid individuals with freebayes, only using sites detected during the previous SNP calling and a ploidy of three. Clonal lineages were determined separately for diploid and triploid as follows: after filtering out sites with a minor allele frequency (MAF) inferior to 5%, more than 50% missing data, and selecting a random SNP for each genomic window of 100 kb, individuals belonging to the same clonal lineage were grouped together with the mlg.filter function in the R package poppr 2.9.3 [80], on the basis of pairwise Euclidean distances with the farthest neighbor method. The cut-off values were chosen visually as suggested previously [81]. The number of clonal lineages was compared between diploid and triploid individuals after correcting sample size by rarefaction, using the R package vegan 2.6-4 [82]. To check whether the proportion of triploid individuals in each site was correlated with latitude, longitude and elevation, we fitted a linear regression model using these three variables in R.

We investigated the population genetic structure of cultivated and wild enset and the origin of triploid individuals using multiple approaches. We obtained a clone-corrected dataset by keeping the sample with the least missing data in each clonal lineage, with the exception of three cultivated individuals which were grouped with wild samples and retained for further analyses. Sites with a MAF < 5% were discarded, and, for each genomic window of 100 kb we kept the site with the higher MAF. Relatedness between clonal lineages was assessed using a method of moment estimator designed for mixed ploidy data [83,84] and implemented in PolyRelatedness v1.11b. Population genetic structure in the clone-corrected dataset was assessed with principal component analysis (PCA) based on individual allele frequencies. F_ST_ values between our three groups (cultivated diploids, cultivated triploids, and wild individuals) were computed with the R package StAMPP 1.6.3 [85], and their 95% confidence intervals (CI) were determined with 1,000 bootstraps across loci.

Genotype likelihoods for each SNP were computed with updog 2.1.2 [86,87] in R, from the read counts in freebayes outputs and with default parameters. Genotype likelihoods were then used to investigate population structure for the entire clone-corrected dataset with entropy [88], a Structure-like algorithm developed for mixed ploidy data. The software was run with a number of populations K ranging from 2 to 15, with five MCMC chains (5,000 burn-in steps, 10,000 MCMC steps, thinning of 10) for each K value to assess convergence. The fit of the data to each value of K was calculated using the Watanabe-Akaike information criterion (WAIC) [89].

### Analysis of phenotypic data for diploid and triploid cultivated individuals

To assess the potential effect of ploidy on enset phenotype, here the pseudostem volume, we used a Bayesian linear mixed-effects approach implemented in the R package brms 2.19.0 [90,91]. First, the pseudostem volume for each individual was calculated according to the shape of a truncated cone, using the measured pseudostem height and basal and apical circumferences. To reduce dimensionality and avoid correlated variables, the environmental data was transformed using PCA. Plant age, ploidy and the two first principal components of the PCA were used as fixed effects, as well as interactions between ploidy, age, and each of the two principal components. The response variable (pseudostem volume) and age values were log-transformed. Membership to clonal lineages was set as a random effect, with the covariance between lineages computed from pairwise relatedness coefficients, to take into account correlation between clonal lineages. Individuals with missing data for any variable and four one-year-old individuals were discarded, leaving us with 555 samples. We fitted the model with four chains of 10,000 iterations, including 2,500 warm-up iterations, using default priors. Convergence was assessed visually by plotting the traces of the chains, and by checking the R-hat values and the effective sample sizes (Bulk-ESS and Tail-ESS). Model fit was evaluated using posterior predictive checks.

## Acknowledgments

We thank the numerous farmers, agricultural extension agents and drivers who generously facilitated our fieldwork. We also thank Romain Pigeault for his help with the mixed-effects model analysis. This research was supported by the Global Challenges Research Fund Agrisystems awards entitled ‘Landscape scale genomic-environment data to enhance the food security of Ethiopian agri-systems’ (Grant No. BB/S014896/1) and ‘Modelling and genomics resources to enhance exploitation of the sustainable and diverse Ethiopian starch crop enset and support livelihoods’ (Grant No. BB/P02307X/1). This research utilised Queen Mary’s Apocrita HPC facility, supported by QMUL Research-IT (http://doi.org/10.5281/zenodo.438045).

## Data Availability

The enset genome assembly and its associated data have been deposited in GenBank BioProject PRJNA899044, and is also available on the Banana Genome Hub website (https://banana-genome-hub.southgreen.fr/). Raw reads from resequencing individuals have been deposited in GenBank BioProject PRJNA1277766, and other data have been deposited in Zenodo (DOI: 10.5281/zenodo.14866251). Scripts used for analysis are available from https://github.com/YannDussert/enset_triploidy_analyses.

## References

1. Heslop-Harrison JS (Pat), Schwarzacher T, Liu Q. Polyploidy: its consequences and enabling role in plant diversification and evolution. Ann Bot. 2023;131: 1–10. doi:10.1093/aob/mcac132

2. Landis JB, Soltis DE, Li Z, Marx HE, Barker MS, Tank DC, et al. Impact of whole-genome duplication events on diversification rates in angiosperms. Am J Bot. 2018;105: 348–363. doi:10.1002/ajb2.1060

3. Leebens-Mack JH, Barker MS, Carpenter EJ, Deyholos MK, Gitzendanner MA, Graham SW, et al. One thousand plant transcriptomes and the phylogenomics of green plants. Nature. 2019;574: 679–685. doi:10.1038/s41586-019-1693-2

4. Salman-Minkov A, Sabath N, Mayrose I. Whole-genome duplication as a key factor in crop domestication. Nat Plants. 2016;2: 1–4. doi:10.1038/nplants.2016.115

5. Akagi T, Jung K, Masuda K, Shimizu KK. Polyploidy before and after domestication of crop species. Curr Opin Plant Biol. 2022;69: 102255. doi:10.1016/j.pbi.2022.102255

6. Otto SP. The evolutionary consequences of polyploidy. Cell. 2007;131: 452–462. doi:10.1016/j.cell.2007.10.022

7. te Beest M, Le Roux JJ, Richardson DM, Brysting AK, Suda J, Kubešová M, et al. The more the better? The role of polyploidy in facilitating plant invasions. Ann Bot. 2012;109: 19–45. doi:10.1093/aob/mcr277

8. Van de Peer Y, Ashman T-L, Soltis PS, Soltis DE. Polyploidy: an evolutionary and ecological force in stressful times. Plant Cell. 2021;33: 11–26. doi:10.1093/plcell/koaa015

9. Meyer RS, DuVal AE, Jensen HR. Patterns and processes in crop domestication: an historical review and quantitative analysis of 203 global food crops. New Phytol. 2012;196: 29–48. doi:10.1111/j.1469-8137.2012.04253.x

10. Borrell JS, Biswas MK, Goodwin M, Blomme G, Schwarzacher T, Heslop-Harrison JS (Pat), et al. Enset in Ethiopia: a poorly characterized but resilient starch staple. Ann Bot. 2019;123: 747–766. doi:10.1093/aob/mcy214

11. Cheesman EE. Classification of the bananas: the genus Ensete Horan. Kew Bull. 1947;2: 97–106. doi:10.2307/4109206

12. Borrell JS, Goodwin M, Blomme G, Jacobsen K, Wendawek AM, Gashu D, et al. Enset-based agricultural systems in Ethiopia: A systematic review of production trends, agronomy, processing and the wider food security applications of a neglected banana relative. Plants People Planet. 2020;2: 212–228. doi:10.1002/ppp3.10084

13. Brandt SA, Spring A, Hiebsch C, McCabe JT, Tabogie E, Diro M, et al. Tree against hunger: enset-based agricultural systems in Ethiopia. American Association for the Advancement of science; 1997.

14. Demissew S, Friis I. Review of the history, taxonomy and nomenclature of ensete and the objectives and expectations of the international workshop on Ensete ventricosum (Welw.) Cheesman. Ethiop J Biol Sci. 2018;17: 1–23.

15. White OW, Biswas MK, Abebe WM, Dussert Y, Kebede F, Nichols RA, et al. Maintenance and expansion of genetic and trait variation following domestication in a clonal crop. Mol Ecol. 2023;32: 4165–4180. doi:10.1111/mec.17033

16. Haile AT, Kovi MR, Johnsen SS, Tesfaye B, Hvoslef-Eide T, Rognli OA. Genetic diversity, population structure and selection signatures in Enset (Ensete ventricosum, (Welw.) Cheesman), an underutilized and key food security crop in Ethiopia. Genet Resour Crop Evol. 2024;71: 1159–1176. doi:10.1007/s10722-023-01683-9

17. Tesfamicael KG, Gebre E, March TJ, Sznajder B, Mather DE, Rodríguez López CM. Accumulation of mutations in genes associated with sexual reproduction contributed to the domestication of a vegetatively propagated staple crop, enset. Hortic Res. 2020;7: 185. doi:10.1038/s41438-020-00409-7

18. Venkatesan L, Muzemil S, Fiche F, Grant M, Studholme DJ. Genome resources for Ensete ventricosum (Enset) and related species. In: Chapman MA, editor. Underutilised Crop Genomes. Cham: Springer International Publishing; 2022. pp. 355–371. doi:10.1007/978-3-031-00848-1_19

19. Yemataw Z, Muzemil S, Ambachew D, Tripathi L, Tesfaye K, Chala A, et al. Genome sequence data from 17 accessions of Ensete ventricosum, a staple food crop for millions in Ethiopia. Data Brief. 2018;18: 285–293. doi:10.1016/j.dib.2018.03.026

20. Birmeta G, Nybom H, Bekele E. Distinction between wild and cultivated enset (Ensete ventricosum) gene pools in Ethiopia using RAPD markers. Hereditas. 2004;140: 139– 148. doi:10.1111/j.1601-5223.2004.01792.x

21. Olango TM, Tesfaye B, Pagnotta MA, Pè ME, Catellani M. Development of SSR markers and genetic diversity analysis in enset (Ensete ventricosum (Welw.) Cheesman), an orphan food security crop from Southern Ethiopia. BMC Genet. 2015;16: 98. doi:10.1186/s12863-015-0250-8

22. Tsegaye A, Struik PC. Influence of repetitive transplanting and leaf pruning on dry matter and food production of enset (Ensete ventricosum Welw. (Cheesman)). Field Crops Res. 2000;68: 61–74. doi:10.1016/S0378-4290(00)00111-8

23. Sharif BM, Burgarella C, Cormier F, Mournet P, Causse S, Van KN, et al. Genome-wide genotyping elucidates the geographical diversification and dispersal of the polyploid and clonally propagated yam (Dioscorea alata). Ann Bot. 2020;126: 1029– 1038. doi:10.1093/aob/mcaa122

24. Sugihara Y, Kudoh A, Oli MT, Takagi H, Natsume S, Shimizu M, et al. Population genomics of yams: evolution and domestication of Dioscorea species. Cham: Springer International Publishing; 2021. pp. 1–28. doi:10.1007/13836_2021_94

25. Chaïr H, Traore RE, Duval MF, Rivallan R, Mukherjee A, Aboagye LM, et al. Genetic diversification and dispersal of taro (Colocasia esculenta (L.) Schott). PLOS ONE. 2016;11: e0157712. doi:10.1371/journal.pone.0157712

26. Howard NP, Micheletti D, Luby JJ, Durel C-E, Denancé C, Muranty H, et al. Pedigree reconstruction for triploid apple cultivars using single nucleotide polymorphism array data. Plants People Planet. 2023;5: 98–111. doi:10.1002/ppp3.10313

27. Van Drunen WE, Husband BC. Evolutionary associations between polyploidy, clonal reproduction, and perenniality in the angiosperms. New Phytol. 2019;224: 1266–1277. doi:10.1111/nph.15999

28. Levin DA. Polyploidy and novelty in flowering plants. Am Nat. 1983;122: 1–25. doi:10.1086/284115

29. Clo J, Kolář F. Short- and long-term consequences of genome doubling: a meta-analysis. Am J Bot. 2021;108: 2315–2322. doi:10.1002/ajb2.1759

30. Mellisse BT, Descheemaeker K, Mourik MJ, Ven GWJ van de. Allometric equations for yield predictions of enset (Ensete ventricosum) and khat (Catha edulis) grown in home gardens of southern Ethiopia. Ann Appl Biol. 2017;171: 95–102. doi:10.1111/aab.12350

31. Negash M, Starr M, Kanninen M. Allometric equations for biomass estimation of Enset (Ensete ventricosum) grown in indigenous agroforestry systems in the Rift Valley escarpment of southern-eastern Ethiopia. Agrofor Syst. 2013;87: 571–581. doi:10.1007/s10457-012-9577-6

32. Lourkisti R, Oustric J, Quilichini Y, Froelicher Y, Herbette S, Morillon R, et al. Improved response of triploid citrus varieties to water deficit is related to anatomical and cytological properties. Plant Physiol Biochem. 2021;162: 762–775. doi:10.1016/j.plaphy.2021.03.041

33. Pacey EK, Maherali H, Husband BC. Polyploidy increases storage but decreases structural stability in Arabidopsis thaliana. Curr Biol. 2022;32: 4057-4063.e3. doi:10.1016/j.cub.2022.07.019

34. Corneillie S, De Storme N, Van Acker R, Fangel JU, De Bruyne M, De Rycke R, et al. Polyploidy affects plant growth and alters cell wall composition. Plant Physiol. 2019;179: 74–87. doi:10.1104/pp.18.00967

35. Guignard MS, Nichols RA, Knell RJ, Macdonald A, Romila C-A, Trimmer M, et al. Genome size and ploidy influence angiosperm species’ biomass under nitrogen and phosphorus limitation. New Phytol. 2016;210: 1195–1206. doi:10.1111/nph.13881

36. Fort A, Ryder P, McKeown PC, Wijnen C, Aarts MG, Sulpice R, et al. Disaggregating polyploidy, parental genome dosage and hybridity contributions to heterosis in Arabidopsis thaliana. New Phytol. 2016;209: 590–599. doi:10.1111/nph.13650

37. Yao H, Dogra Gray A, Auger DL, Birchler JA. Genomic dosage effects on heterosis in triploid maize. Proc Natl Acad Sci. 2013;110: 2665–2669. doi:10.1073/pnas.1221966110

38. De Storme N, Copenhaver GP, Geelen D. Production of diploid male gametes in Arabidopsis by cold-induced destabilization of postmeiotic radial microtubule arrays. Plant Physiol. 2012;160: 1808–1826. doi:10.1104/pp.112.208611

39. Pécrix Y, Rallo G, Folzer H, Cigna M, Gudin S, Le Bris M. Polyploidization mechanisms: temperature environment can induce diploid gamete formation in Rosa sp. J Exp Bot. 2011;62: 3587–3597. doi:10.1093/jxb/err052

40. Drenth A, Kema G. The vulnerability of bananas to globally emerging disease threats. Phytopathology. 2021;111: 2146–2161. doi:10.1094/PHYTO-07-20-0311-RVW

41. Henry IM, Dilkes BP, Young K, Watson B, Wu H, Comai L. Aneuploidy and genetic variation in the Arabidopsis thaliana triploid response. Genetics. 2005;170: 1979–1988. doi:10.1534/genetics.104.037788

42. Ramsey J, Schemske DW. Pathways, mechanisms, and rates of polyploid formation in flowering plants. Annual Review of Ecology and Systematics. 1998. pp. 467–501.

43. Adeleke MTV, Pillay M, Okoli BE. Relationships between meiotic irregularities and fertility in diploid and triploid Musa L. Cytologia (Tokyo). 2004;69: 387–393. doi:10.1508/cytologia.69.387

44. Park SM, Wakana A, Kim JH, Jeong CS. Male and female fertility in triploid grapes (Vitis complex) with special reference to the production of aneuploid plants. Vitis. 2002;41: 11–20.

45. Tamrat S, Borrell JS, Shiferaw E, Wondimu T, Kallow S, Davies RM, et al. Reproductive biology of wild and domesticated Ensete ventricosum: Further evidence for maintenance of sexual reproductive capacity in a vegetatively propagated perennial crop. Plant Biol. 2022;24: 482–491. doi:10.1111/plb.13390

46. Cheng H, Concepcion GT, Feng X, Zhang H, Li H. Haplotype-resolved de novo assembly using phased assembly graphs with hifiasm. Nat Methods. 2021;18: 170– 175. doi:10.1038/s41592-020-01056-5

47. Laetsch DR, Blaxter ML. BlobTools: Interrogation of genome assemblies. F1000Research. 2017;6: 1287.

48. Guan D, McCarthy SA, Wood J, Howe K, Wang Y, Durbin R. Identifying and removing haplotypic duplication in primary genome assemblies. Bioinformatics. 2020;36: 2896– 2898. doi:10.1093/bioinformatics/btaa025

49. Putnam NH, O’Connell BL, Stites JC, Rice BJ, Blanchette M, Calef R, et al. Chromosome-scale shotgun assembly using an in vitro method for long-range linkage. Genome Res. 2016;26: 342–350. doi:10.1101/gr.193474.115

50. Li H, Durbin R. Fast and accurate short read alignment with Burrows-Wheeler transform. Bioinformatics. 2009;25: 1754–1760. doi:10.1093/bioinformatics/btp324

51. Dierckxsens N, Mardulyn P, Smits G. NOVOPlasty: de novo assembly of organelle genomes from whole genome data. Nucleic Acids Res. 2017;45: e18. doi:10.1093/nar/gkw955

52. Camacho C, Coulouris G, Avagyan V, Ma N, Papadopoulos J, Bealer K, et al. BLAST+: Architecture and applications. BMC Bioinformatics. 2009;10: 421–421. doi:10.1186/1471-2105-10-421

53. Quinlan AR, Hall IM. BEDTools: a flexible suite of utilities for comparing genomic features. Bioinformatics. 2010;26: 841–842. doi:10.1093/bioinformatics/btq033

54. Wu C-S, Sudianto E, Chiu H-L, Chao C-P, Chaw S-M. Reassessing banana phylogeny and organelle inheritance modes using genome skimming data. Front Plant Sci. 2021;12. doi:10.3389/fpls.2021.713216

55. Flynn JM, Hubley R, Goubert C, Rosen J, Clark AG, Feschotte C, et al. RepeatModeler2 for automated genomic discovery of transposable element families. Proc Natl Acad Sci. 2020;117: 9451–9457. doi:10.1073/pnas.1921046117

56. Smit AFA, Hubley R, Green P. RepeatMasker Open-4.0. 2013–2015. 2015.

57. Wang Z, Rouard M, Biswas MK, Droc G, Cui D, Roux N, et al. A chromosome-level reference genome of Ensete glaucum gives insight into diversity and chromosomal and repetitive sequence evolution in the Musaceae. GigaScience. 2022;11: giac027. doi:10.1093/gigascience/giac027

58. D’Hont A, Denoeud F, Aury J-M, Baurens F-C, Carreel F, Garsmeur O, et al. The banana (Musa acuminata) genome and the evolution of monocotyledonous plants. Nature. 2012;488: 213–217. doi:10.1038/nature11241

59. Hao Z, Lv D, Ge Y, Shi J, Weijers D, Yu G, et al. RIdeogram: drawing SVG graphics to visualize and map genome-wide data on the idiograms. PeerJ Comput Sci. 2020;6: e251. doi:10.7717/peerj-cs.251

60. Bolger AM, Lohse M, Usadel B. Trimmomatic: a flexible trimmer for Illumina sequence data. Bioinformatics. 2014;30: 2114–2120. doi:10.1093/bioinformatics/btu170

61. Dobin A, Davis CA, Schlesinger F, Drenkow J, Zaleski C, Jha S, et al. STAR: ultrafast universal RNA-seq aligner. Bioinformatics. 2013;29: 15–21. doi:10.1093/bioinformatics/bts635

62. Li H, Handsaker B, Wysoker A, Fennell T, Ruan J, Homer N, et al. The sequence alignment/map format and SAMtools. Bioinformatics. 2009;25: 2078–2079. doi:10.1093/bioinformatics/btp352

63. Kriventseva EV, Kuznetsov D, Tegenfeldt F, Manni M, Dias R, Simão FA, et al. OrthoDB v10: sampling the diversity of animal, plant, fungal, protist, bacterial and viral genomes for evolutionary and functional annotations of orthologs. Nucleic Acids Res. 2019;47: D807–D811. doi:10.1093/nar/gky1053

64. Droc G, Martin G, Guignon V, Summo M, Sempéré G, Durant E, et al. The banana genome hub: a community database for genomics in the Musaceae. Hortic Res. 2022;9: uhac221. doi:10.1093/hr/uhac221

65. Brůna T, Hoff KJ, Lomsadze A, Stanke M, Borodovsky M. BRAKER2: automatic eukaryotic genome annotation with GeneMark-EP+ and AUGUSTUS supported by a protein database. NAR Genomics Bioinforma. 2021;3: qaa108. doi:10.1093/nargab/lqaa108

66. Hoff KJ, Lange S, Lomsadze A, Borodovsky M, Stanke M. BRAKER1: Unsupervised RNA-Seq-Based Genome Annotation with GeneMark-ET and AUGUSTUS. Bioinformatics. 2016;32: 767–769. doi:10.1093/bioinformatics/btv661

67. Mapleson D, Garcia Accinelli G, Kettleborough G, Wright J, Clavijo BJ. KAT: a K-mer analysis toolkit to quality control NGS datasets and genome assemblies. Bioinformatics. 2017;33: 574–576. doi:10.1093/bioinformatics/btw663

68. Ranallo-Benavidez TR, Jaron KS, Schatz MC. GenomeScope 2.0 and Smudgeplot for reference-free profiling of polyploid genomes. Nat Commun. 2020;11: 1432. doi:10.1038/s41467-020-14998-3

69. Manni M, Berkeley MR, Seppey M, Simão FA, Zdobnov EM. BUSCO Update: novel and streamlined workflows along with broader and deeper phylogenetic coverage for scoring of eukaryotic, prokaryotic, and viral genomes. Mol Biol Evol. 2021;38: 4647– 4654. doi:10.1093/molbev/msab199

70. Karger D, Conrad O, Böhner J, Kawohl T, Kreft H, Soria-Auza RW, et al. Climatologies at high resolution for the earth’s land surface areas. Sci Data. 2017;4: 170122. doi:10.1038/sdata.2017.122

71. Karger D, Conrad O, Böhner J, Kawohl T, Kreft H, Soria-Auza R, et al. Data from: Climatologies at high resolution for the earth’s land surface areas. Dryad Digit Repos. 2017. doi:10.5061/dryad.kd1d4

72. Miki Y, Yoshida K, Enoki H, Komura S, Suzuki K, Inamori M, et al. GRAS-Di system facilitates high-density genetic map construction and QTL identification in recombinant inbred lines of the wheat progenitor Aegilops tauschii. Sci Rep. 2020;10: 21455. doi:10.1038/s41598-020-78589-4

73. Garrison E, Marth G. Haplotype-based variant detection from short-read sequencing. arXiv. 2012 [cited 25 July 2022]. doi:10.48550/arXiv.1207.3907

74. Danecek P, Auton A, Abecasis G, Albers CA, Banks E, DePristo MA, et al. The variant call format and VCFtools. Bioinformatics. 2011;27: 2156–2158. doi:10.1093/bioinformatics/btr330

75. Knaus BJ, Grünwald NJ. vcfr: a package to manipulate and visualize variant call format data in R. Mol Ecol Resour. 2017;17: 44–53. doi:10.1111/1755-0998.12549

76. Poncet P. modeest: Mode estimation. 2019. Available: https://cran.r-project.org/web/packages/modeest/index.html

77. Marçais G, Kingsford C. A fast, lock-free approach for efficient parallel counting of occurrences of k-mers. Bioinformatics. 2011;27: 764–770. doi:10.1093/bioinformatics/btr011

78. Zarrei M, Wilkin P, Ingrouille MJ, Leitch IJ, Buerki S, Fay MF, et al. Speciation and evolution in the Gagea reticulata species complex (Tulipeae; Liliaceae). Mol Phylogenet Evol. 2012;62: 624–639. doi:10.1016/j.ympev.2011.11.003

79. Meirmans PG, Liu S, van Tienderen PH. The analysis of polyploid genetic data. J Hered. 2018;109: 283–296. doi:10.1093/jhered/esy006

80. Kamvar ZN, Tabima JF, Grünwald NJ. Poppr: An R package for genetic analysis of populations with clonal, partially clonal, and/or sexual reproduction. PeerJ. 2014;2014: 1–14. doi:10.7717/peerj.281

81. Arnaud-Haond S, Duarte CM, Alberto F, Serrão EA. Standardizing methods to address clonality in population studies. Mol Ecol. 2007;16: 5115–5139. doi:10.1111/j.1365-294X.2007.03535.x

82. Oksanen J, Simpson GL, Blanchet FG, Kindt R, Legendre P, Minchin PR, et al. vegan: Community ecology package. 2022. Available: https://cran.r-project.org/web/packages/vegan/index.html

83. Huang K, Ritland K, Guo S, Shattuck M, Li B. A pairwise relatedness estimator for polyploids. Mol Ecol Resour. 2014;14: 734–744. doi:10.1111/1755-0998.12217

84. Huang K, Ritland K, Guo S, Dunn DW, Chen D, Ren Y, et al. Estimating pairwise relatedness between individuals with different levels of ploidy. Mol Ecol Resour. 2015;15: 772–784. doi:10.1111/1755-0998.12351

85. Pembleton LW, Cogan NOI, Forster JW. StAMPP: an R package for calculation of genetic differentiation and structure of mixed-ploidy level populations. Mol Ecol Resour. 2013;13: 946–952. doi:10.1111/1755-0998.12129

86. Gerard D, Ferrão LFV, Garcia AAF, Stephens M. Genotyping polyploids from messy sequencing data. Genetics. 2018;210: 789–807. doi:10.1534/genetics.118.301468

87. Gerard D, Ferrão LFV. Priors for genotyping polyploids. Bioinformatics. 2020;36: 1795– 1800. doi:10.1093/bioinformatics/btz852

88. Shastry V, Adams PE, Lindtke D, Mandeville EG, Parchman TL, Gompert Z, et al. Model-based genotype and ancestry estimation for potential hybrids with mixed-ploidy. Mol Ecol Resour. 2021;21: 1434–1451. doi:10.1111/1755-0998.13330

89. Watanabe S. Asymptotic aquivalence of Bayes cross validation and widely applicable information criterion in singular learning theory. J Mach Learn Res. 2010;11: 3571– 3594.

90. Bürkner P-C. brms: an R package for Bayesian multilevel models using Stan. J Stat Softw. 2017;80: 1–28. doi:10.18637/jss.v080.i01

91. Carpenter B, Gelman A, Hoffman MD, Lee D, Goodrich B, Betancourt M, et al. Stan: a probabilistic programming language. J Stat Softw. 2017;76: 1–32. doi:10.18637/jss.v076.i01

