## Supplementary tables and figures for "Recurrent evolution of cryptic triploids in cultivated enset increases yield"

**Supplementary Table 1 Statistics of the *Ensete ventricosum* genome assembly**

| | | |
|---|---|---|
| | EnVent_Maze_1.0 (this study) | Bedadeti_annotated (Yemataw et al. 2018) |
| *Assembly* | | |
| Assembly size (Mb) | 534.05 | 451.28 |
| Number of scaffolds | 806 | 47,742 |
| N50 / L50 (size in kb / count) | 57,407.2 / 5 | 21.1 / 6,010 |
| N90 / L90 (size in kb / count) | 35,910.5 / 9 | 4.0 / 23,334 |
| Largest scaffold (Mb) | 68.01 | 0.21 |
| % GC | 39.8 | 38.9 |
| % of repeat elements | 58.6 | - |
| BUSCO genome | C:98.6% [S:93.7%, D:4.9%], F:0.4% | C:84.2% [S:80.7%, D:3.5%], F:10.8% |
| *Gene annotation* | | |
| Number of protein-coding genes | 35,238 | 58,438 |
| BUSCO proteome | C:96.2% [S:91.5%, D:4.7%], F:1.7% | C:25.4% [S:24.3%, D:1.1%], F:22.4% |

BUSCO analyses have been carried out with the embryophyta_odb10 data set. C: complete genes, S: single copy, D: duplicated, F: fragmented genes.

**Supplementary Table 2 Result summary of the Bayesian linear mixed-effects analysis**

| | Estimate | Est. error | l-95% CI | u-95% CI | Rhat | Bulk_ESS | Tail_ESS |
|---|---|---|---|---|---|---|---|
| *Group-Level Effects* | | | | | | | |
| sd(Intercept) | 1.36 | 0.32 | 0.74 | 2.02 | 1.00 | 5242 | 6046 |
| sd(logAge) | 0.81 | 0.21 | 0.40 | 1.24 | 1.00 | 4737 | 4782 |
| cor(Intercept,logAge) | -0.95 | 0.03 | -0.98 | -0.78 | 1.00 | 6720 | 6759 |
| *Population-Level Effects* | | | | | | | |
| Intercept | 9.41 | 0.51 | 8.35 | 10.45 | 1.00 | 16578 | 18301 |
| logAge | 1.68 | 0.31 | 1.03 | 2.33 | 1.00 | 16699 | 18837 |
| pc1 | -0.10 | 0.06 | -0.22 | 0.03 | 1.00 | 16407 | 21673 |
| ploidy3n | -1.70 | 0.85 | -3.54 | -0.11 | 1.00 | 11609 | 15461 |
| pc2 | 0.51 | 0.10 | 0.30 | 0.71 | 1.00 | 16608 | 19874 |
| logAge:pc1 | 0.05 | 0.04 | -0.03 | 0.12 | 1.00 | 16794 | 21542 |
| logAge:ploidy3n | 1.21 | 0.52 | 0.22 | 2.32 | 1.00 | 12060 | 15965 |
| pc1:ploidy3n | 0.08 | 0.15 | -0.21 | 0.37 | 1.00 | 13008 | 17117 |
| logAge:pc2 | -0.33 | 0.07 | -0.46 | -0.19 | 1.00 | 16967 | 20557 |
| ploidy3n:pc2 | -0.47 | 0.24 | -0.95 | 0.00 | 1.00 | 11381 | 17030 |
| logAge:pc1:ploidy3n | 0.01 | 0.09 | -0.17 | 0.19 | 1.00 | 13677 | 18959 |
| logAge:ploidy3n:pc2 | 0.28 | 0.15 | -0.01 | 0.59 | 1.00 | 12000 | 17536 |
| *Family Specific Parameters* | | | | | | | |
| sigma | 0.62 | 0.02 | 0.58 | 0.67 | 1.00 | 13669 | 18172 |

Estimate: median value of the posterior distribution. Est. error: median absolute deviation

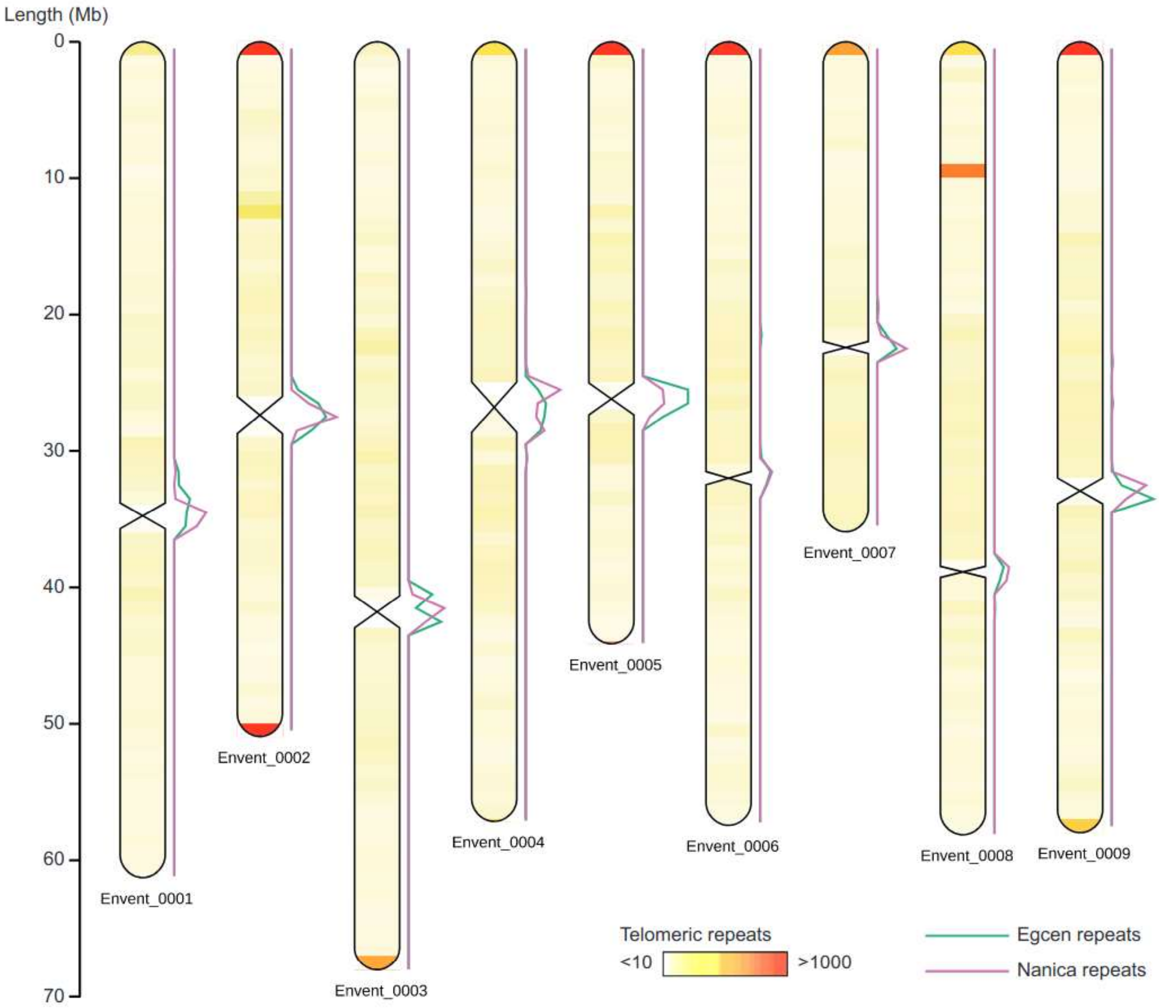

**Supplementary Figure 1 Chromosome-scale assembly of the *Ensete ventricosum* genome.** The nine largest scaffolds are represented, with constriction points at the likely positions of centromeres. The number of canonical telomeric repeats is indicated by a color gradient inside each scaffold ideogram, and the relative abundance of Egcen and Nanica repeats, which are found in centromeric regions in other Musaceae, by colored lines on the right side of each ideogram. The number of all repeat types have been computed for 1 Mb windows along chromosomes. The maximum number of Egcen copies per window has been capped at 250, to allow for better readability.

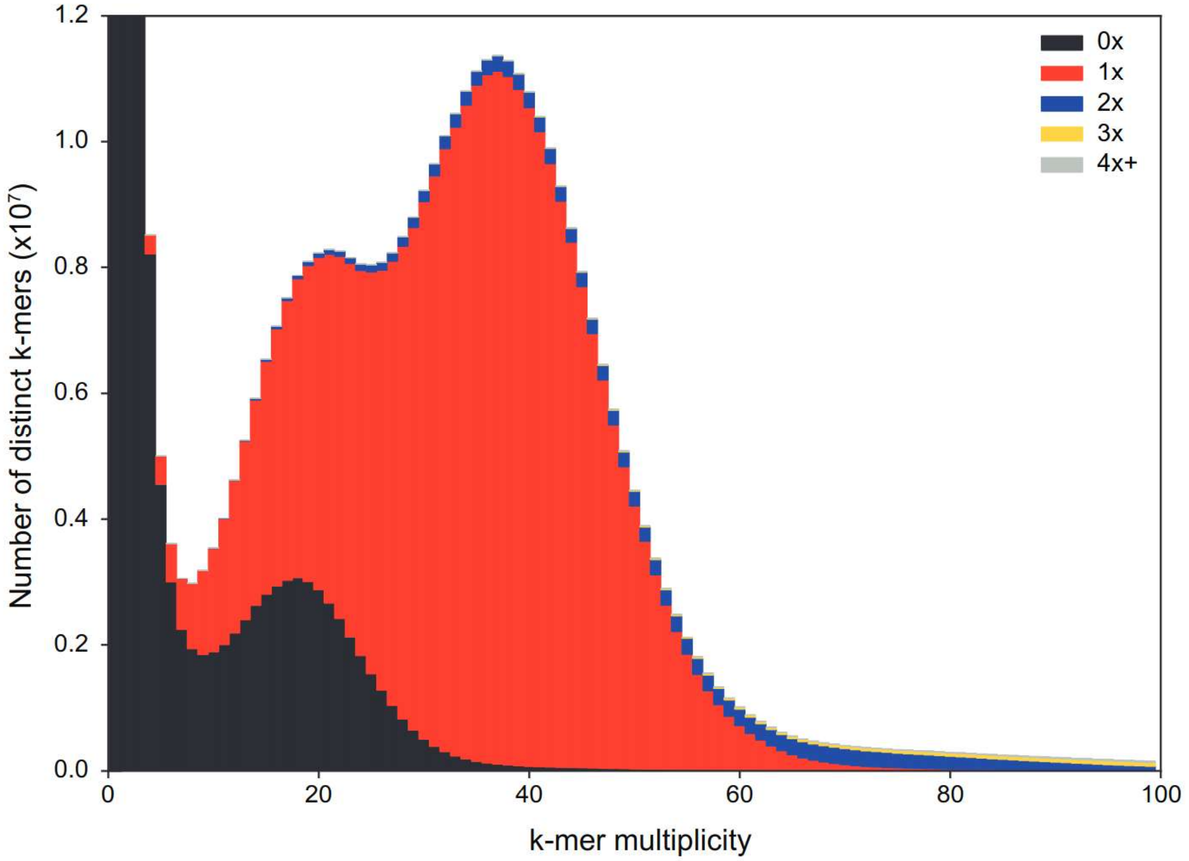

**Supplementary Figure 2 K-mer content comparison between the CCS PacBio reads and the *Ensete ventricosum* genome assembly.** The k-mer spectrum (k = 21) for the PacBio reads is represented as a stacked histogram, with colored areas corresponding to k-mers absent from the assembly (0x), k-mers occurring once in the assembly (1x), k-mers occuring twice (2x), etc. Peaks at multiplicities 21 and 37 represent the heterozygous and homozygous content in the reads, respectively. The low proportion of homozygous k-mers found twice in the assembly (in blue) and the proportion of absent k-mer (in black) in the heterozygous peak are indicative of a good pseudo-haploid assembly.

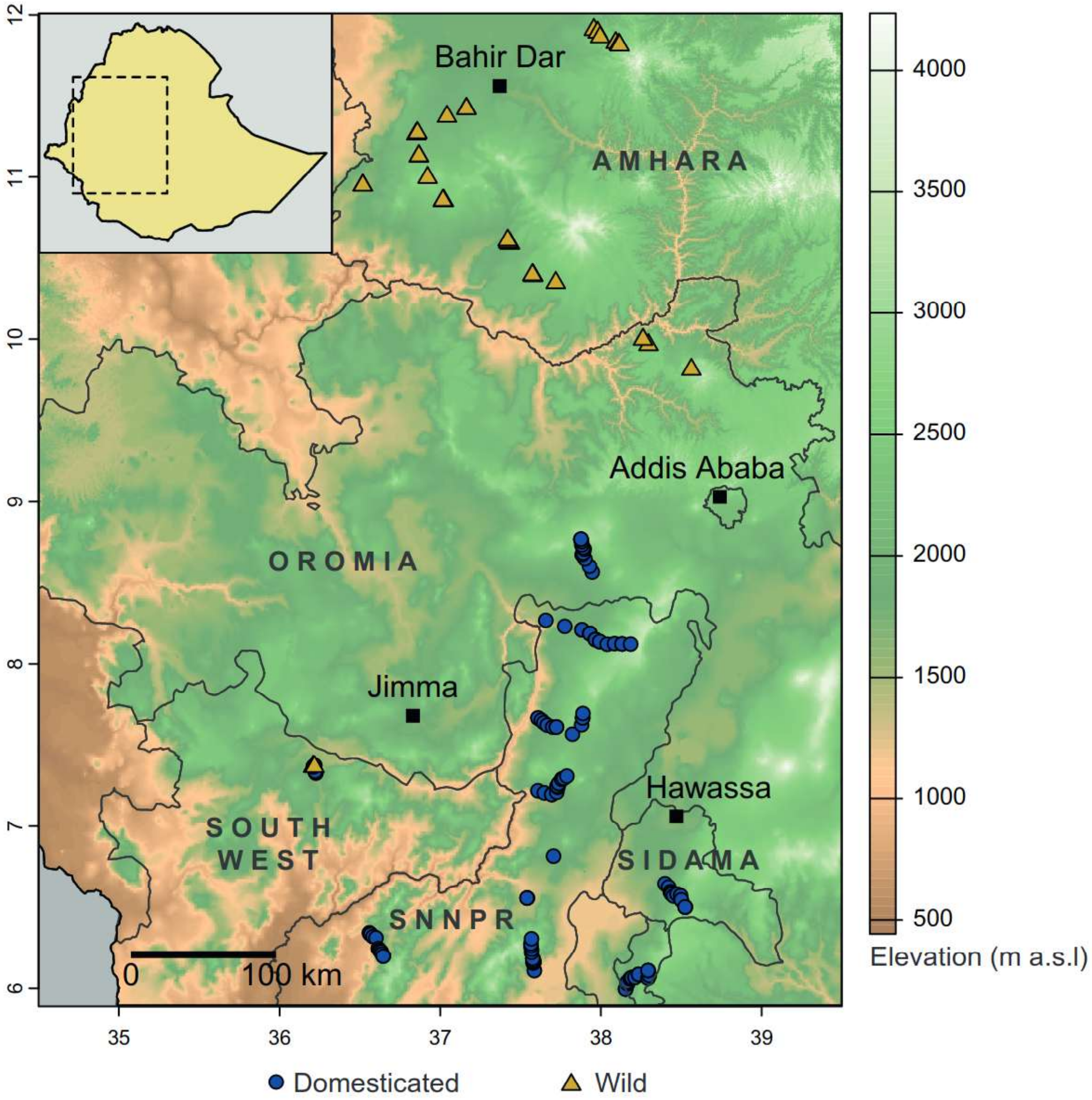

**Supplementary Figure 3 Geographical distribution of the 658 cultivated and 65 wild individuals of *Ensete ventricosum* sampled in Ethiopia**. Cultivated individuals are represented by blue circles, wild ones by yellow triangles. Light gray lines delimit regional states, with their names indicated in bold. Major cities are represented by black squares. The inset at top left hand corner shows the total area of Ethiopia, with the dashed square indicating the location of the enlarged map.

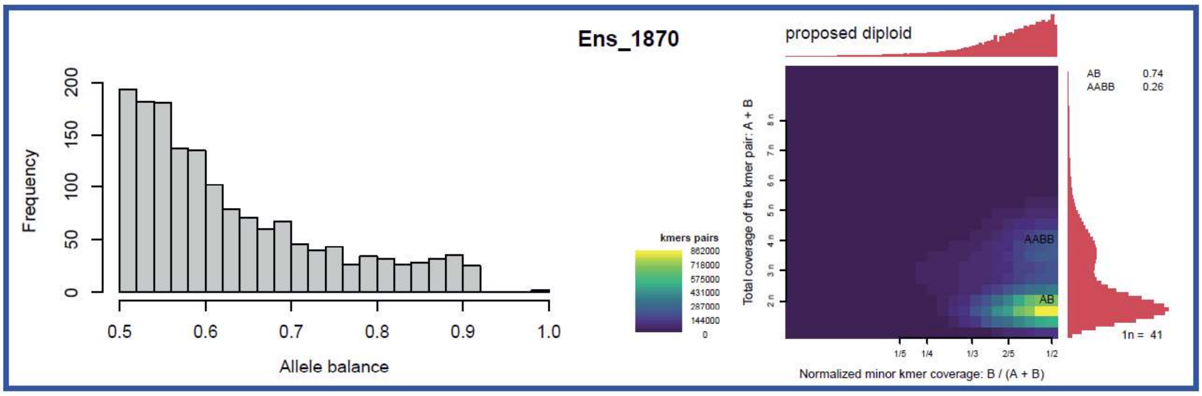
Ens_1870
Frequency
Allele balance
kmers pairs
862000
718000
575000
431000
287000
144000
0
proposed diploid
Total coverage of the kmer pair: A + B
Normalized minor kmer coverage: B / (A + B)
AABB
AB
AB 0.74
AABB 0.26
1n = 41

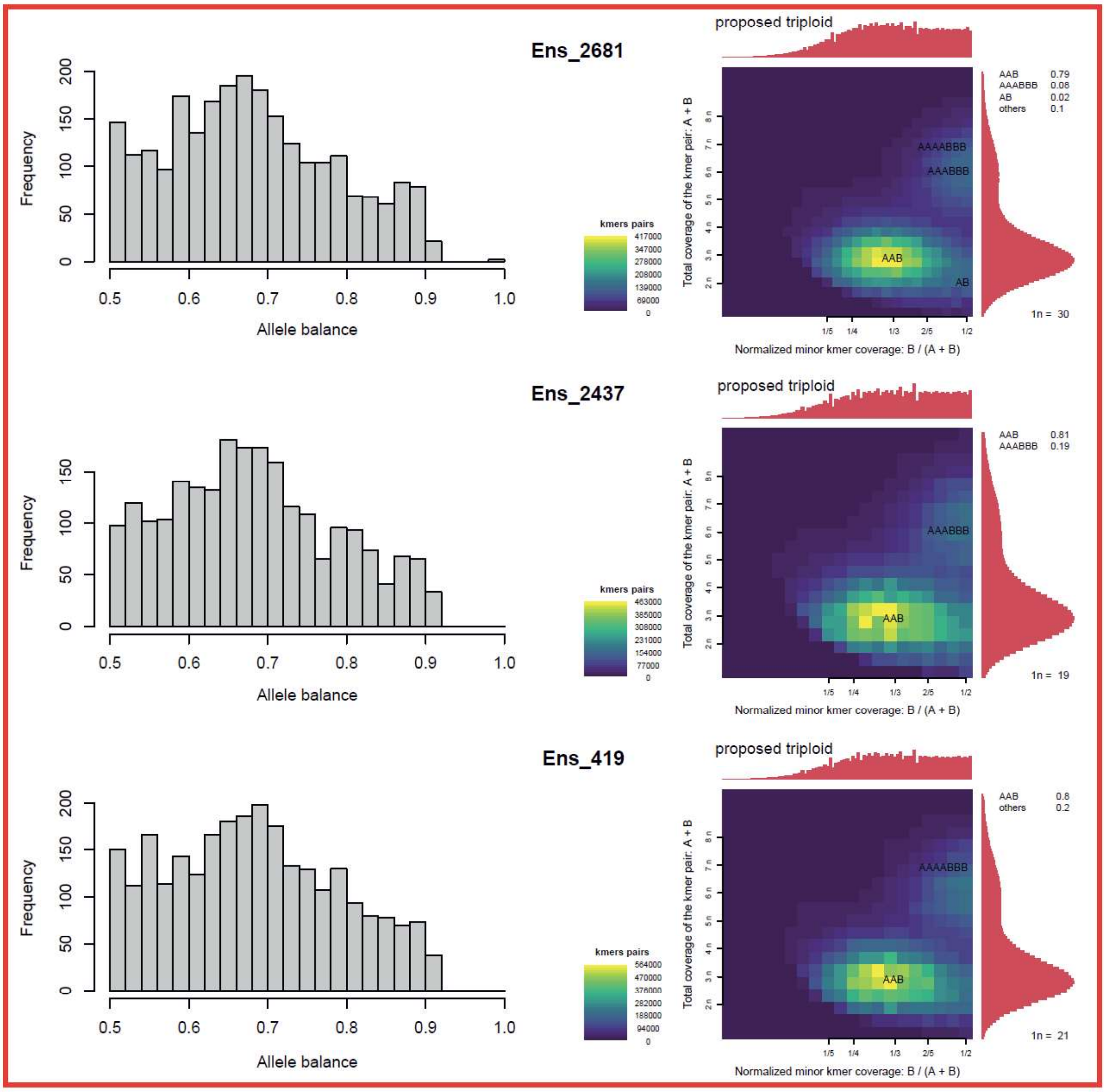
Ens_2681
Frequency
Allele balance
kmers pairs
417000
347000
278000
208000
139000
69000
0
proposed triploid
Total coverage of the kmer pair: A + B
Normalized minor kmer coverage: B / (A + B)
AAAABBB
AAABBB
AAB
AB
AAB 0.79
AAABBB 0.08
AB 0.02
others 0.1
1n = 30
Ens_2437
Frequency
Allele balance
kmers pairs
463000
385000
308000
231000
154000
77000
0
proposed triploid
Total coverage of the kmer pair: A + B
Normalized minor kmer coverage: B / (A + B)
AAABBB
AAB
AAB 0.81
AAABBB 0.19
1n = 19
Ens_419
Frequency
Allele balance
kmers pairs
564000
470000
376000
282000
188000
94000
0
proposed triploid
Total coverage of the kmer pair: A + B
Normalized minor kmer coverage: B / (A + B)
AAAABBB
AAB
AAB 0.8
others 0.2
1n = 21

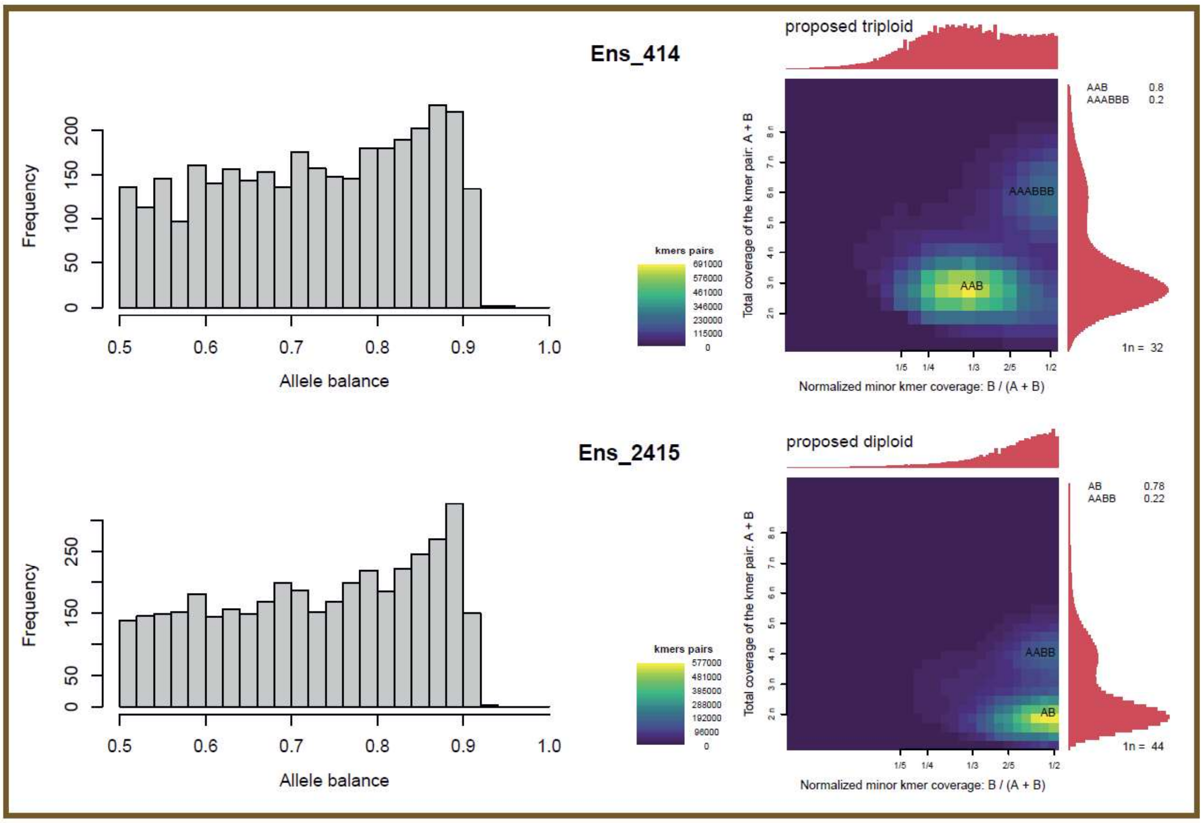

**Supplementary Figure 4 Allele balance distributions and SmudgePlots for resequenced enset samples.** For each resequenced individual (rows), the allele balance distribution from the GRAS-Di data (left panels) is compared to the SmudgePlot k-mer analysis from the whole-genome resequencing data (right panels). In SmudgePlots, heatmap colors indicate the number of heterozygous (ie, differing by only one nucleotide) 21-mer pairs in each bin, from 0 in dark purple to the maximum value in bright yellow. Histograms represent the total coverage of k-mer pairs for each axis. The brighter "smudges" indicate the main ploidy of the sample, i.e. AAB for triploids or AB for diploids. Colored boxes around individuals indicate whether they were classified in the diploid (red), triploid (blue) or aberrant (brown) group on the basis of their allele balance distribution.

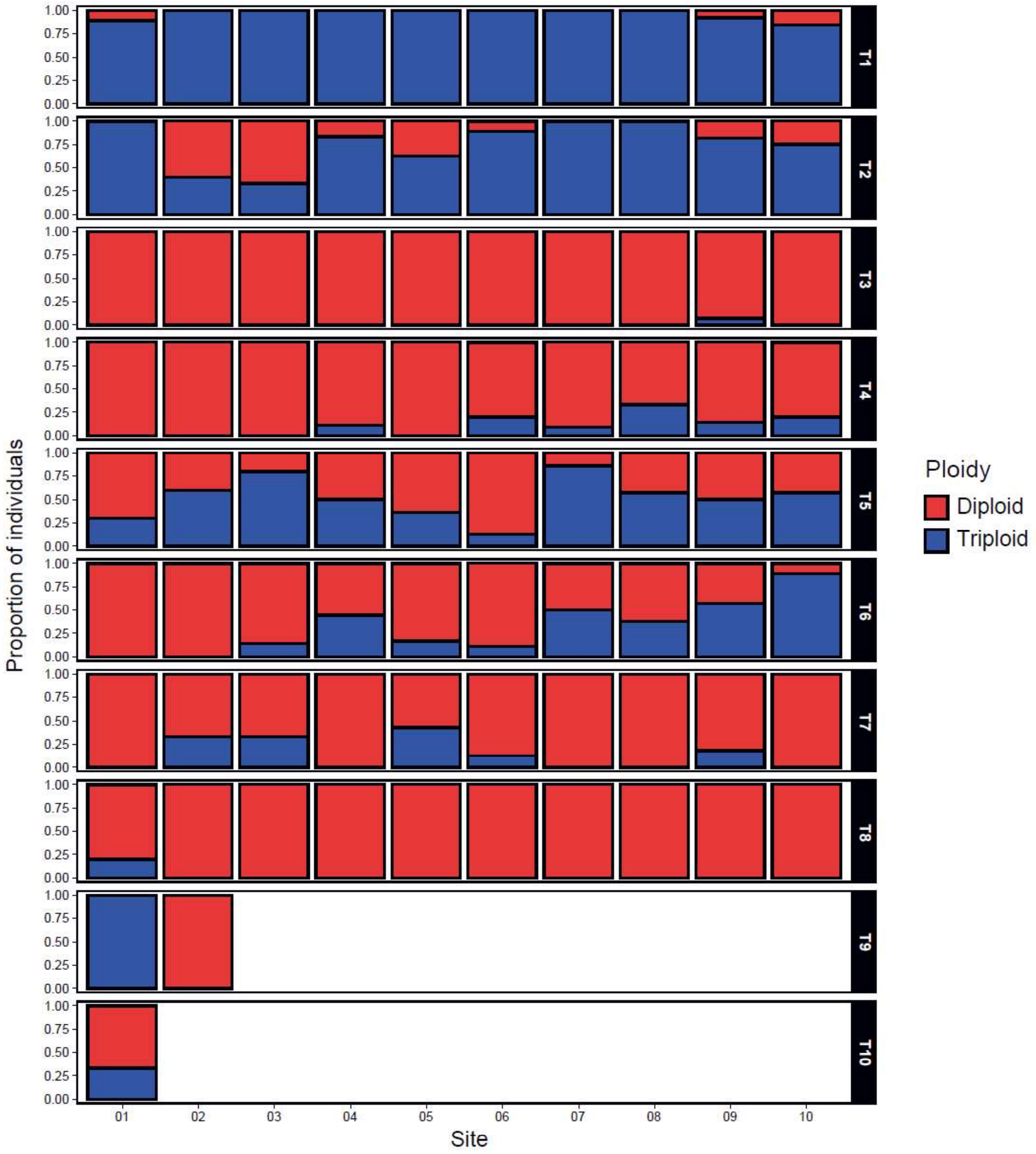

**Supplementary Figure 5 Ploidy variation in cultivated enset in southwestern Ethiopia.** The proportion of individuals with a diploid or triploid cytotype is represented by a stacked barplot for each site (on the x axis) in each transect (in each panel). Sites within transects are ordered by altitude, with the lowest altitude to the left.

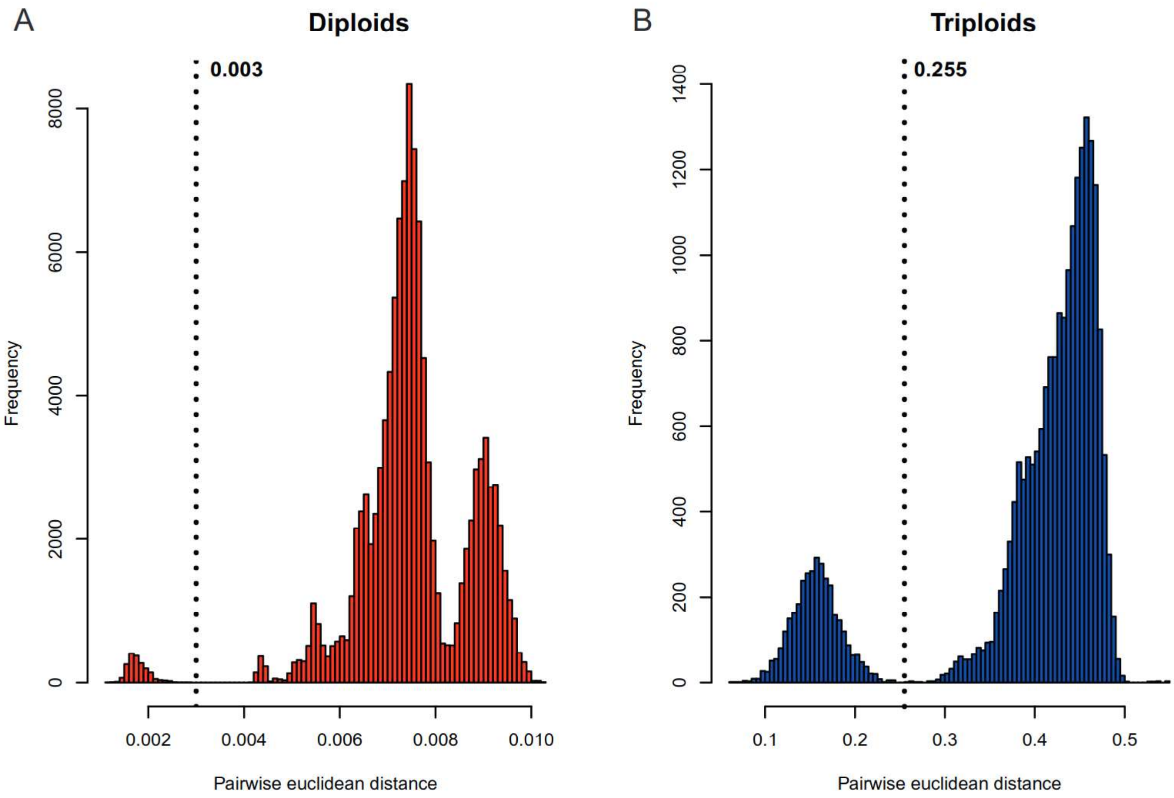

**Supplementary Figure 6 Distribution of pairwise Euclidean genetic distances in enset, between diploid (A) and triploid (B) individuals.** The dotted vertical line represents the distance threshold used to group samples into clonal lineages for each ploidy level, as the primary peaks correspond to pairs of samples with a very similar multi-locus genotype.

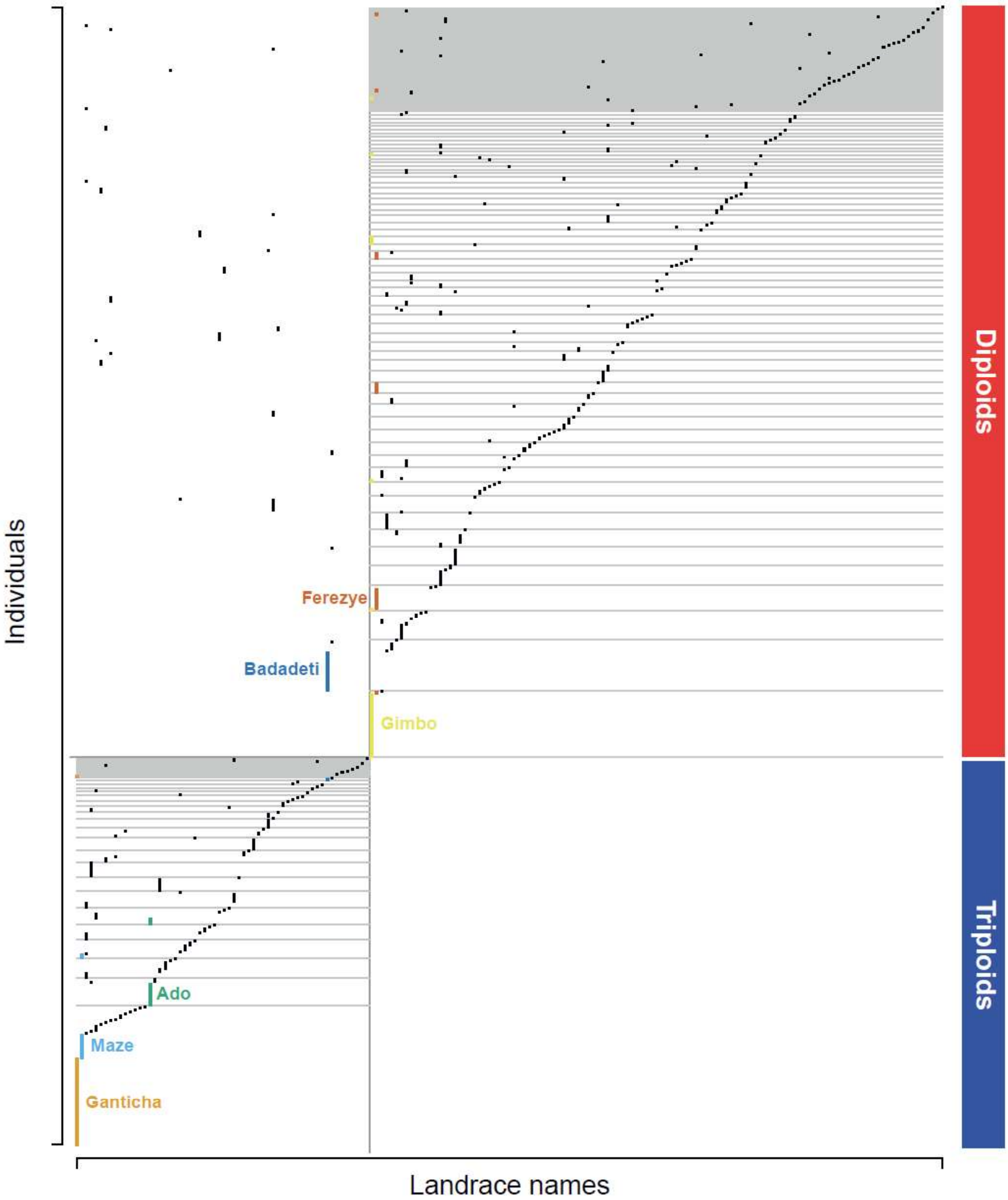

**Supplementary Figure 7 Relationship between vernacular landrace names and clonal lineages for domesticated *Ensete ventricosum*.** Landrace names are on the x-axis and individuals on the y-axis. Clonal lineages are separated by vertical light gray lines on the left for triploids and on the right for diploids. For clarity, only the six most common varieties have been annotated with their name, and highlighted with a color matching their label.

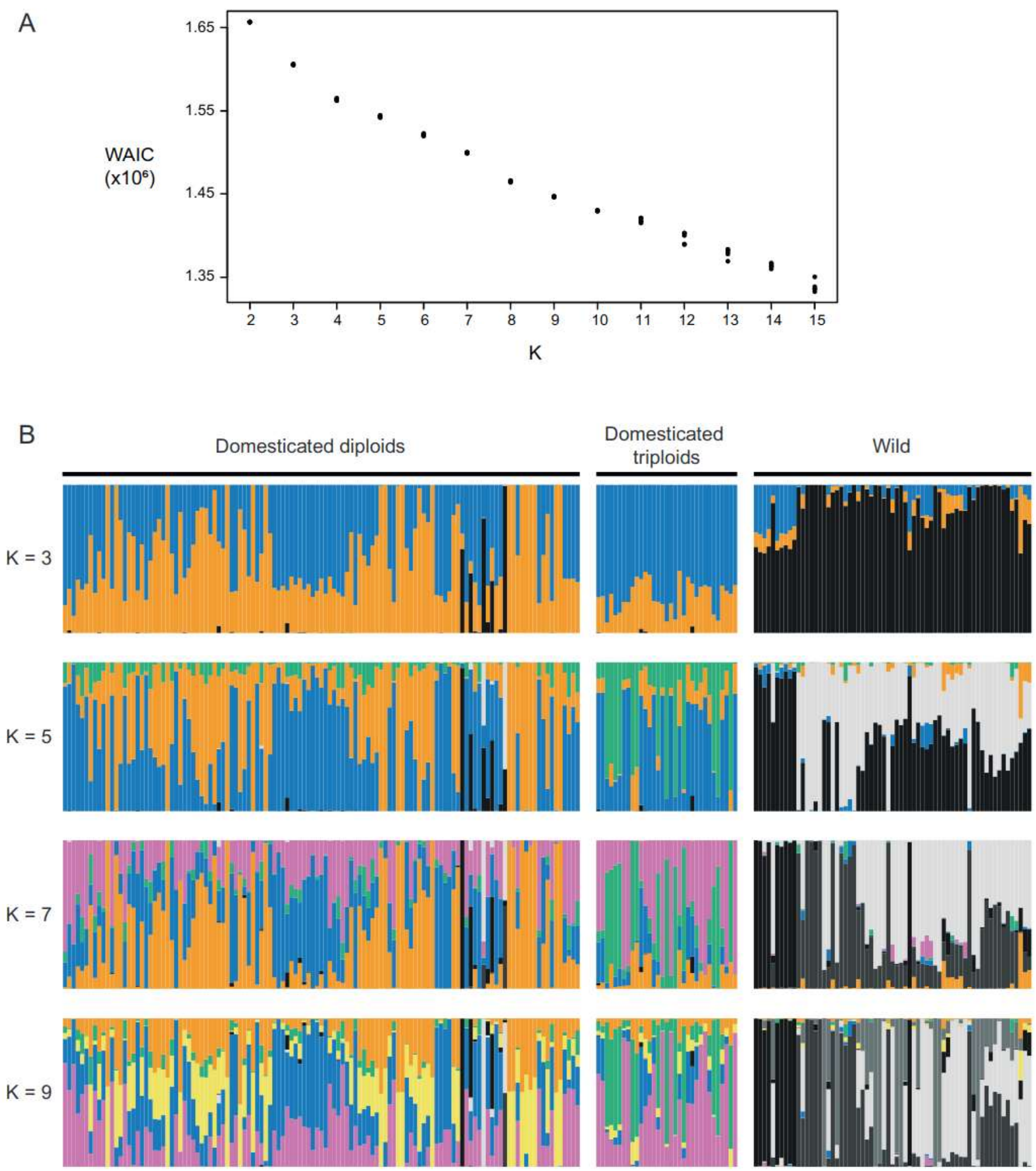

**Supplementary Figure 8 Genetic structure of cultivated and wild enset, based on the entropy algorithm.** A: Values of the Watanabe-Akaike information criterion (WAIC) for each number K of populations. B: Individual membership coefficients in each genetic group for different values of K. Each individual is represented as a vertical bar, with colored segments corresponding to the estimated membership coefficients in each cluster.
